# Yeast DNA Polymerase δ Proofreading Nuclease Suppresses Translesion Synthesis and Facilitates Recombination in DNA Lesion Bypass

**DOI:** 10.1101/2025.01.14.632785

**Authors:** Jun Che, Quan Wang, Jiapeng He, Zhao Zhang, Jee Min Chung, Danielle N. Gallagher, Xin Hong, Hai Rao, Eun Shim, James E. Haber, Hengyao Niu, Sang Eun Lee

## Abstract

Translesion synthesis (TLS) employs specialized polymerases to incorporate nucleotides opposite DNA lesions, allowing replication to continue temporarily without repairing the lesions, while sacrificing replication fidelity (*1*). TLS depends on a ubiquitin molecule attached to the clamp PCNA at residue Lys^164^ (*2, 3*); the precise biochemical and molecular basis how PCNA modification stimulates TLS is still elusive (*4-10*). In this study, we discovered that DNA polymerase δ (Pol δ) 3’ to 5’ “proofreading” exonuclease degrades nascent TLS synthesis opposite pyrimidine dimers and inhibits TLS on both leading and lagging strands. Rad18 E3 ligase and PCNA ubiquitylation become dispensable for TLS when Pol δ exonuclease is inactive. Biochemical reconstitution showed that the major UV lesion bypass polymerase Rad30 is intrinsically inefficient at bypassing thymidine dimers and requires multiple turnovers. Pol δ exonuclease impedes Rad30 activity both catalytically and non-catalytically during and post-lesion bypass. Importantly, PCNA and PCNA ubiquitylation enhance processivity of Rad30 after thymidine dimer bypass, a critical step for further DNA synthesis extension and overcoming Pol δ inhibition. Our study reveals a new paradigm how PCNA ubiquitylation facilitates TLS and the new role of Pol δ nuclease in lesion bypass pathway choice and replication fidelity at fork-blocking DNA lesions.

## Main text

Complete and accurate genome duplication is essential for cell division and chromosomal integrity. However, DNA replication is often interrupted by DNA lesions, unique DNA structures, and tightly bound proteins, all of which can halt replication fork progression (*11, 12*). Stalled replication fork should bypass an obstacle or restart downstream to the lesion to avoid replication fork collapse and DNA double strand break, which can lead to chromosome aberrations and rearrangements, a hallmark of cancer cells.

TLS is the evolutionary conserved lesion bypass mechanism and is a primary inducer of damage-induced mutagenesis. Current model posits that UV or MMS-induced TLS in yeast requires multiple rounds of polymerase switching: the switch from the replicative polymerases Pol ε or Pol δ to one or more TLS polymerases upon encountering a DNA lesion, and then switch back to reinstate Pol ε or Pol δ for bulk DNA replication after lesion bypass (**Fig. 1A**). Since Pol ε and Pol δ also possess 3’ to 5’ proofreading exonuclease activity that removes unpaired nucleotides, we hypothesize that upon the second switch, reinstated Pol ε or Pol δ nuclease might recognize nucleotides opposite UV or MMS lesions as replication errors, trim them, and repress TLS (**Fig. 1A**). To test this idea, we analyzed TLS activity and UV-induced mutagenesis in *pol2-04, pol3-01*, or *pol3-5DV* mutants (*13, 14*), which are defective in either Pol ε (Pol2) or Pol δ (Pol3) proofreading nuclease activity, respectively, using several genetic assays. First, we deleted the key recombination genes *RAD51* or *RAD52* from *pol2* or *pol3* mutants and examined their viability after UV or MMS damage. Deletion of *RAD51* or *RAD52* makes TLS the primary means for UV/MMS tolerance. If proofreading nucleases antagonize TLS, then nuclease-deficient mutations would confer UV or MMS resistance in *rad51*Δ or *rad52*Δ cells due to an increased TLS. Interestingly, inactivation of Pol ε exonuclease activity (*pol2-04*) does not provide UV resistance in *rad51*Δ or *rad52*Δ mutants (**Fig. S1**). In contrast, *pol3-5DV* or *pol3-01* suppresses UV sensitivity in *rad51*Δ or *rad52*Δ cells (**Fig. 1B, Fig. S2A**). Increased UV resistance depends on all three TLS polymerases (Rev1, Rev3, and Rad30; **Fig. 1C** and **Fig. S2 B-D**). The nuclease-deficient mutants with intact *RAD51* and *RAD52* show only mild UV resistance and do not exhibit UV sensitivity in *rev1 rev3 rad30* (*tls*Δ) cells, where recombination is the primary lesion bypass option (**Fig. S2 E**-**F**). The effect of Pol δ exonuclease defects on cell viability is lesion-type specific: *pol3-5DV* and *pol3-01* mutations do not affect survival after HU or IR treatments (**Fig. S3 A-B**). The *pol3-5DV* mutant also did not affect ectopic homologous recombinational repair (*15*) after inducing a site-specific DSB in *rad51*Δ cells (**Fig. S3 C**). The *pol3-5DV* mutation still suppresses UV sensitivity in nucleotide excision repair (NER) deficient *rad14*Δ cells, suggesting that the effect of *pol3-5DV* on the viability of *rad51*Δ cells after UV irradiation is unlikely due to improved UV lesion repair (**Fig. S4**).

**Fig. 1.**
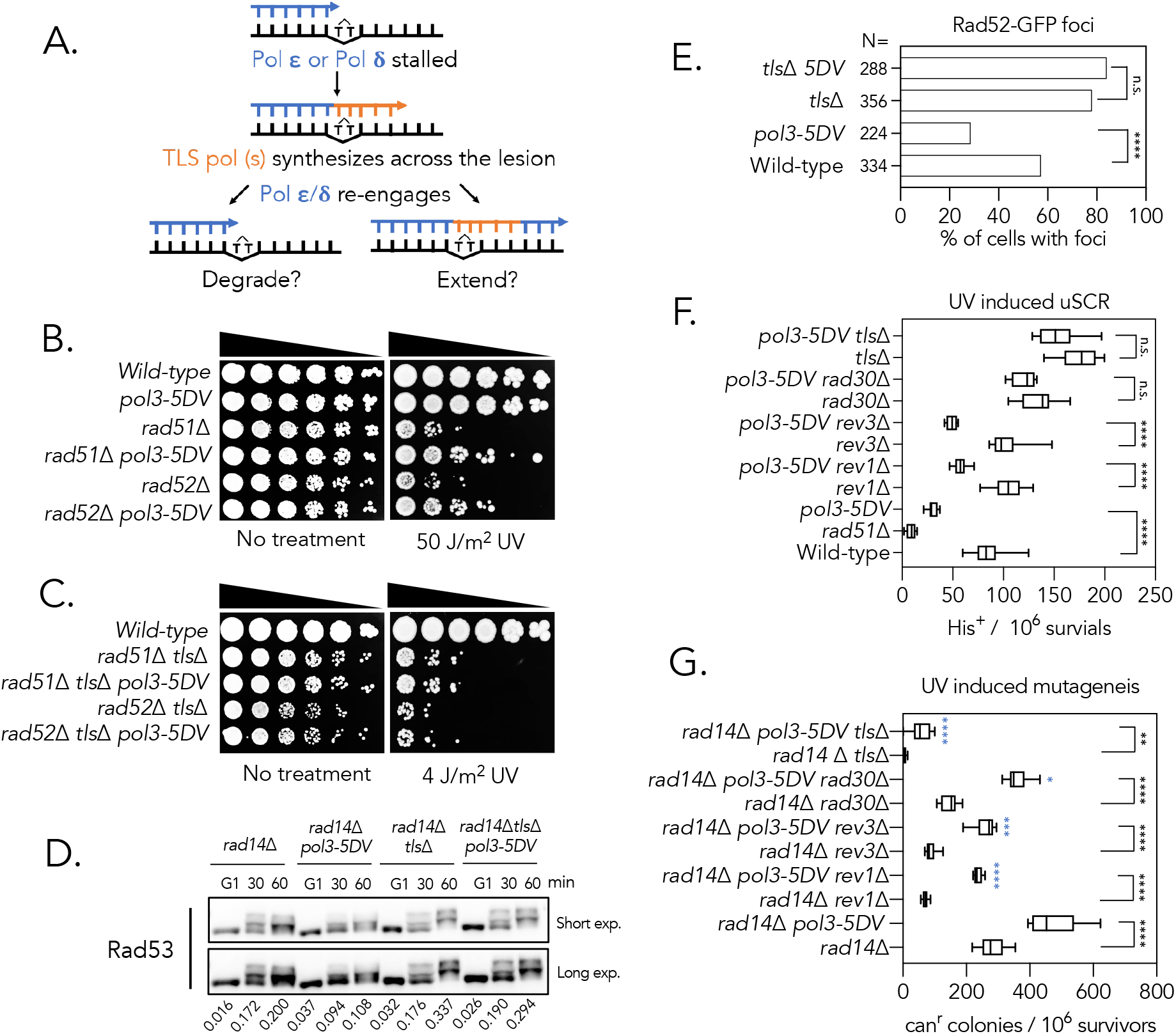
Pol δ proofreading exonuclease suppresses TLS and facilitates recombination. **A**. Schematic diagram of polymerase-switch model of translesion synthesis; degradation or extension of nascent TLS product are two possible options after the replicative polymerase is reinstated; **B-C**. Sensitivity of cells with indicated genotypes to different doses of 254 nm ultraviolet light (UV) treatment. 5-fold serial dilutions of cells were spotted onto YEP-dextrose (YEPD) plates and treated with or without UV. *tls*Δ represents *rev1*Δ *rev3*Δ *rad30*Δ triple mutant; **D**. Western blot detecting Rad53 from the indicated cells. Cells were G_1_ arrested, treated with 1 J/m^2^ UV, released into cell cycle and harvested at different time-points. The percentage of (hyper gel-shifted) phosphorylated Rad53 relative to the total amount of Rad53 in individual lanes were quantified using ImageJ and shown as the level of checkpoint activation; **E**. Percentage of log-phase cells with indicated genotypes showing Rad52-GFP foci after 20 J/m^2^ UV treatment and 2h culture; **F**. Frequency of UV (50 J/m^2^) induced unequal Sister chromatid recombination (uSCR, schematic assay diagram is shown in **Fig. S2G**) in cells with indicated genotypes; **G**. Forward mutation frequency at *CAN1* gene in cells with indicated genotypes after 3 J/m^2^ UV irradiation. Blue stars (****) indicate statistical significance when comparing to the values of *rad14*Δ *pol3-5DV cells. P*-values were calculated by Chi-square-test for **Fig.1E** and two-tailed parametric unpaired t-test for **Fig.1F & 1G**. Statistical significance: \**P*<0.05, \*\**P* <0.01,*** *P*<0.001 **** *P* <0.0001, “n.s.” not significant.

### Pol δ nuclease promotes recombination while suppressing TLS

UV or MMS treatment induces single-strand DNA (ssDNA) gaps at stalled replication forks or nascent daughter strands and triggers DNA damage checkpoint cascades that can be analyzed by detecting Rad53 phosphorylation in yeast. Since TLS plays a crucial role in repairing ssDNA gaps, we predict that elevated TLS in the nuclease mutants would reduce the level of ssDNA gaps and attenuate checkpoint activation. We found that phosphorylated Rad53 is reduced in *rad14*Δ *pol3-5DV* cells in a TLS-dependent manner after UV treatment (**Fig. 1D**). *Pol3-5DV* cells also show reduced UV-induced Rad52-GFP foci formation in a TLS-dependent manner (**Fig. 1E**). By scoring the number of His3^*+*^ colonies in cells bearing *his3* direct repeats (*16*), we also found that the frequency of unequal sister chromatid recombination (uSCR) is decreased in *pol3-5DV* cells after UV irradiation (**Fig. 1F, Fig. S2G** for schematic uSCR assay). Inactivation of TLS effectively offsets the uSCR defect in *pol3-5DV*, indicating that TLS and recombination are competing for ssDNA gap repair and that Pol δ exonuclease promotes recombination while suppressing TLS (**Fig. 1F**).

TLS is prone to induce mutations because of the more relaxed active site architecture of TLS polymerases (*17*). If *pol3-5DV* elevates TLS upon UV treatment, we predict that *pol3-5DV* should also increase UV-induced mutagenesis in a TLS-dependent manner. We measured the forward mutation frequency at the *CAN1* gene in *pol3-5DV rad14*Δ cells upon UV irradiation. Deletion of *RAD14* helps discriminate TLS-associated mutagenesis from that produced by NER. *Pol3-5DV* significantly increases both spontaneous and UV-induced mutagenesis at the *CAN1* locus (**Fig. 1G & Table. S2**). Spontaneous mutagenesis in *pol3-5DV* is not TLS-dependent and is likely attributed to the failure to correct mis-incorporated nucleotides during DNA replication (**Table. S2**). In contrast, UV-induced mutagenesis in *pol3-5DV* depends on all three TLS polymerases (**Fig. 1G**). When all three TLS genes were deleted, the UV-induced mutagenesis in *pol3-5DV* cells was almost indistinguishable to that in *POL3* WT cells (**Fig. 1G**). These results suggest that polymerase δ proofreading exonuclease suppresses TLS in yeast.

### PCNA mono-ubiquitylation is dispensable for TLS in *pol3-5DV* mutants

The choice between TLS and recombination is tightly linked to PCNA K164 ubiquitylation in yeast cells (*18, 19*). Rad6/Rad18 (E2/E3)-dependent PCNA mono-ubiquitylation is required for TLS, while Ubc13/Mms2 (E2)/Rad5 (E3)-dependent PCNA polyubiquitylation drives recombination (*2, 3*). We assessed UV or MMS sensitivity in *ubc13*Δ, *mms2*Δ, *rad5*Δ, or *rad18*Δ cells carrying *pol3-5DV* or *pol3-01* mutations. Both *pol3-5DV* and *pol3-01* mutations suppress UV or MMS sensitivity in *ubc13*Δ, *mms2*Δ, *rad5*Δ or *rad18*Δ mutants (**Fig. 2A, B&D & Fig. S5 A-B**). This suppression depends on all three TLS polymerases, and the mutations act recessively (**Fig. 2C&D & Fig. S5C**). These results suggest that Pol δ proofreading suppresses TLS independently of PCNA mono- or poly-ubiquitylation. In yeast, Siz1 induces PCNA sumoylation and modulates ssDNA gap repair by inhibiting the Rad51-dependent salvage recombination pathway (*20, 21*). Accordingly, *siz1*Δ suppresses UV/MMS sensitivity in *rad5*Δ or *rad18*Δ cells in a Rad51 and Rad52-dependent manner. We found that *pol3-5DV siz1*Δ or *pol3-01 siz1*Δ mutations further suppress UV and MMS sensitivity in *rad5*Δ or *rad18*Δ cells compared to *siz1* or *pol3-5DV* mutations alone (**Fig. 2E & Fig. S5 A-B**). *Pol3-5DV* or *pol3-01* also suppresses UV sensitivity in *ubc13*Δ cells independently of Rad51 (**Fig. S5 D&E**) and in *pcna-K164R* mutants, which abolish PCNA ubiquitylation and most sumoylation (*2*) (**Fig. 2F**). These results suggest that Pol δ proofreading nuclease antagonizes TLS independently of homologous recombination, PCNA K164 ubiquitylation and sumoylation.

**Fig. 2.**
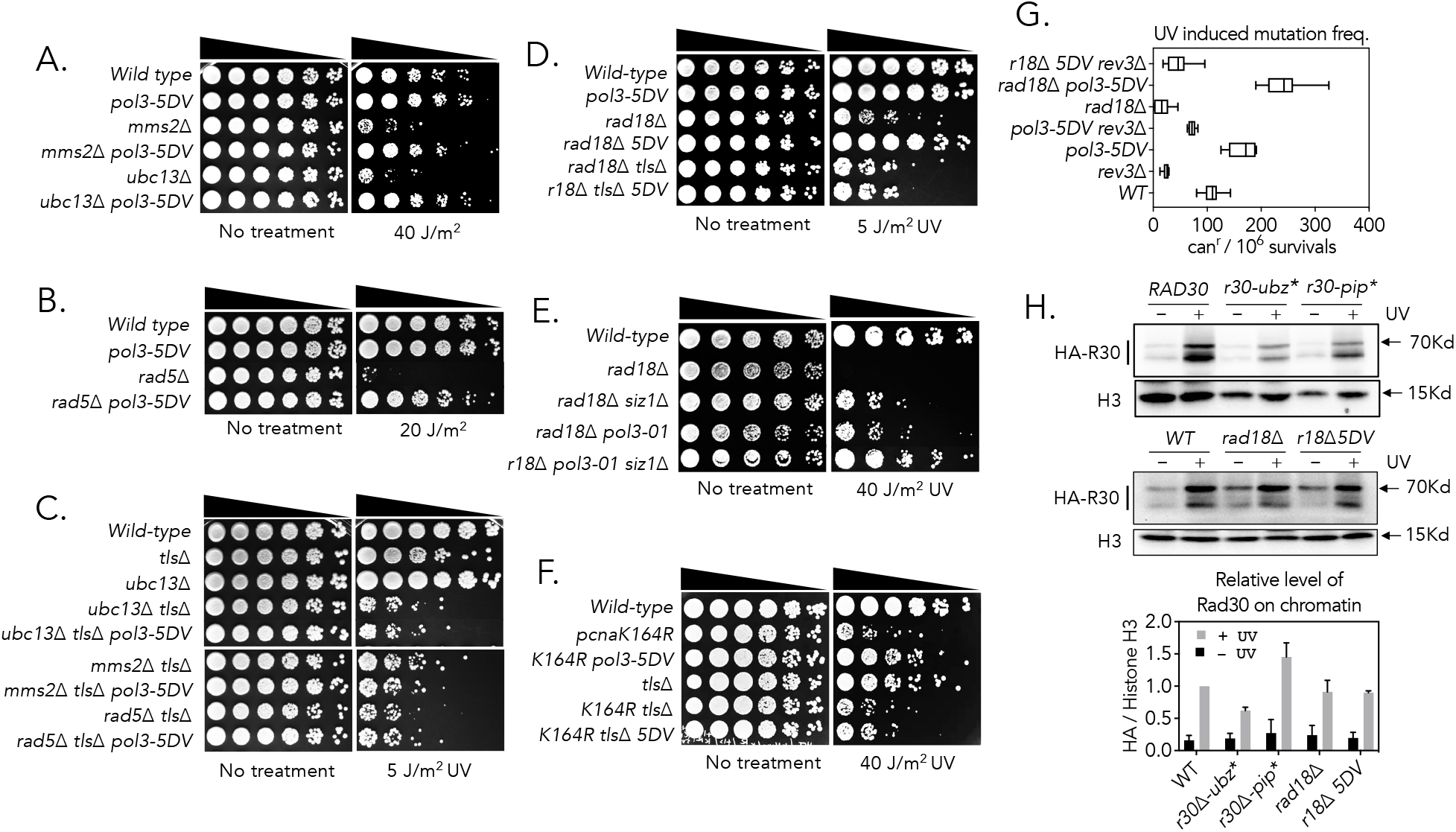
Pol δ exonuclease suppresses TLS independent to PCNA ubiquitylation or sumoylation. **A-F**. Sensitivity of cells to different doses of 254 nm ultraviolet light treatment. 5-fold serial dilutions of cells with indicated genotypes were spotted onto YEPD plates and treated with or without UV; **G**. UV (30 J/m^2^) induced forward mutation frequency at *CAN1* gene in cells with indicated genotypes. **H**. Western blot detecting chromatin bound N-terminal HA-tagged Rad30 or rad30 mutants expressed from its native promoter at genomic locus before and after 100 J/m^2^ UV treatment and the quantification.

We also found that UV-induced *CAN1* mutation frequency in *pol3-5DV rad18*Δ cells was significantly (2.2-fold) elevated than in wild-type cells (**Fig. 2G**). Deletion of *REV3* almost completely abolished the UV-induced mutation frequency in *pol3-5DV rad18*Δ cells (**Fig. 2G**). The results suggest that Rad18 becomes dispensable for TLS in *pol3-5DV* cells.

The prevailing view is that Rad18-dependent PCNA mono-ubiquitylation promotes TLS by recruiting TLS polymerases and/or facilitating the switch from replicative to TLS polymerases via their ubiquitin-binding domains (*4, 22-27*). This raises the question how *pol3-5DV* bypasses the requirement for PCNA mono-ubiquitylation in TLS polymerase recruitment and/or polymerase switch. To address this, we analyzed the effect of *pol3-5DV* on the recruitment of Rad30, the single-subunit TLS polymerase, at DNA lesions. Besides its polymerization domain, Rad30 contains an UBZ (ubiquitin-binding) and a PIP (PCNA-interacting peptide) domains, which are crucial for its TLS function (*23, 28*). Accordingly, *rad30-D570A* (ubz^-^) or *F627AF628A* (pip^-^) mutations render cells sensitive to UV treatment. Interestingly, *pol3-5DV* suppresses UV sensitivity in these mutants (**Fig. S5F**), suggesting that interactions between Rad30 and PCNA or PCNA-ubiquitylation are dispensable for its TLS function in *pol3-5DV* cells.

In yeast, TLS polymerases do not form discrete nuclear foci after UV treatment, and direct evidence of Rad18 facilitating TLS polymerase recruitment to DNA lesions *in vivo* is still lacking. We developed a modified chromatin fractionation assay which detects Rad30 association to the formaldehyde crosslinked chromatin (see Materials and Methods). We found that neither *rad30-D570A* nor *rad30-FFAA* mutation significantly reduce Rad30 recruitment (**Fig. 2H**). Furthermore, the levels of chromatin-bound Rad30 were similar in *rad18*Δ and wild-type cells after UV treatment. The *pol3-5DV* mutation did not improve Rad30 recruitment to chromatin after UV treatment (**Fig. 2H**). These results suggest that neither PCNA ubiquitylation nor the UBZ domain of Rad30 are essential for Rad30 recruitment to chromatin in yeast (**Fig. 2H**). Instead, Rad18 and PCNA ubiquitylation likely play a role in other steps of TLS, beyond Rad30 recruitment.

### Biochemical analysis of thymidine dimer bypass by Rad30

To further explore the mechanism of TLS regulation by Pol δ nuclease, we reconstituted thymidine dimer bypass by Rad30 in biochemical assays. We expressed and purified Rad30 with an N-terminal FLAG tag in yeast cells, along with PCNA produced in bacteria (**Fig. S6A and B**). A ubiquitin-PCNA fusion was constructed to mimic PCNA monoubiquitylation (*23*) and produced in bacteria (**Fig. S6C**). Similarly to previous reports (*28, 29*), under low salt and close to physiological magnesium conditions (50 mM KCl and 2 mM Mg^2+^), Rad30 (20 nM) efficiently bypassed the template T-T dimer on a primer/template substrate (C26). However, raising the salt concentration to 150 mM resulted in significant pausing of Rad30 at the T-T dimer site, with minimal bypass observed (**Fig. S7A**). While titrating Rad30 from 20 nM to 60 nM produced only a modest increase in lesion bypass (from ∼5% to ∼10%), a similar titration under 5 mM Mg^2+^ led to nearly a fivefold increase in T-T dimer bypass (from ∼10% to over 50%) (**Fig. S7B**). Given that Rad30 likely has a higher turnover rate at 5 mM Mg^2+^, we tested lesion bypass in single-turnover reactions by adding heparin following Rad30-substrate complex formation to trap Rad30 that is unbound or dissociated from the DNA substrate (*30*) .In this assay, we used substrates with the priming site positioned immediately before the template thymidine dimer (C28) and a non-damaged template strand as a control. Similarly, as before, under 2 mM Mg^2+^ and 50 mM KCl, Rad30 efficiently bypassed the thymidine dimer (Lane 6), exhibiting polymerase activity comparable to that on non-damaged DNA (Lane 2) (**Fig. S7C**). Increasing the salt to 150 mM KCl again largely abolished Rad30’s lesion bypass activity (Lane 8) (**Fig. S7C**), whereas initiation on the non-damaged substrate remained efficient, albeit with a reduced final product length (Lane 4) (**Fig. S7C**) consistent with the previous observation that the polymerase activity of Rad30 is salt sensitive (*28, 29*). For a single-turnover reaction under 50 mM KCl, we preincubated the substrates with Rad30 in the absence of dNTP, followed by the addition of heparin before introducing dNTP to initiate the polymerase reaction. Under these conditions, 1-2 base pair synthesis was detected on the non-damaged substrate (Lane 3) (**Fig. S7C**), which aligns with the accommodation of two template nucleotides at a time by Rad30/Polη (*31*). However, the presence of heparin largely abolished lesion bypass, even at 50 mM KCl (Lane 7) (**Fig. S7C**), indicating that T-T dimer lesion bypass is likely a multi-turnover reaction.

We next investigated the impact of PCNA on Rad30-catalyzed T-T dimer lesion bypass. Under 5 mM Mg^2+^, both PCNA and ubiquitinylated PCNA promoted the formation of longer extension products, with a more pronounced effect from ubiquitinylated PCNA (**Fig. 3A**), which became even more evident at 200 mM KCl (**Fig. 3A**). Similarly, Rad30 stimulation was also observed under 2 mM Mg^2+^ despite an overall lower reaction rate (**Fig. S8**). These results suggest that PCNA enhances Rad30 processivity during lesion bypass (*28*), which may be further increased by PCNA mono-ubiquitination. In addition to improved processivity, a modest increase in lesion bypass was also noticed in the presence of PCNA or ubiquitinylated PCNA consistent with a previous finding on the abasic site containing substrate (*5, 28*). To confirm this observation, we performed T-T dimer bypass assays using only dATP to trap bypass products under more stringent conditions (200 mM KCl, 30 min reaction). Here, the addition of PCNA or ubiquitinylated PCNA led to nearly a two-fold increase in lesion bypass (**Fig. 3B**). Thus, we conclude that PCNA and ubiquitinylated PCNA have a comparable effect on the stimulation of Rad30 in T-T dimer bypass.

**Fig. 3.**
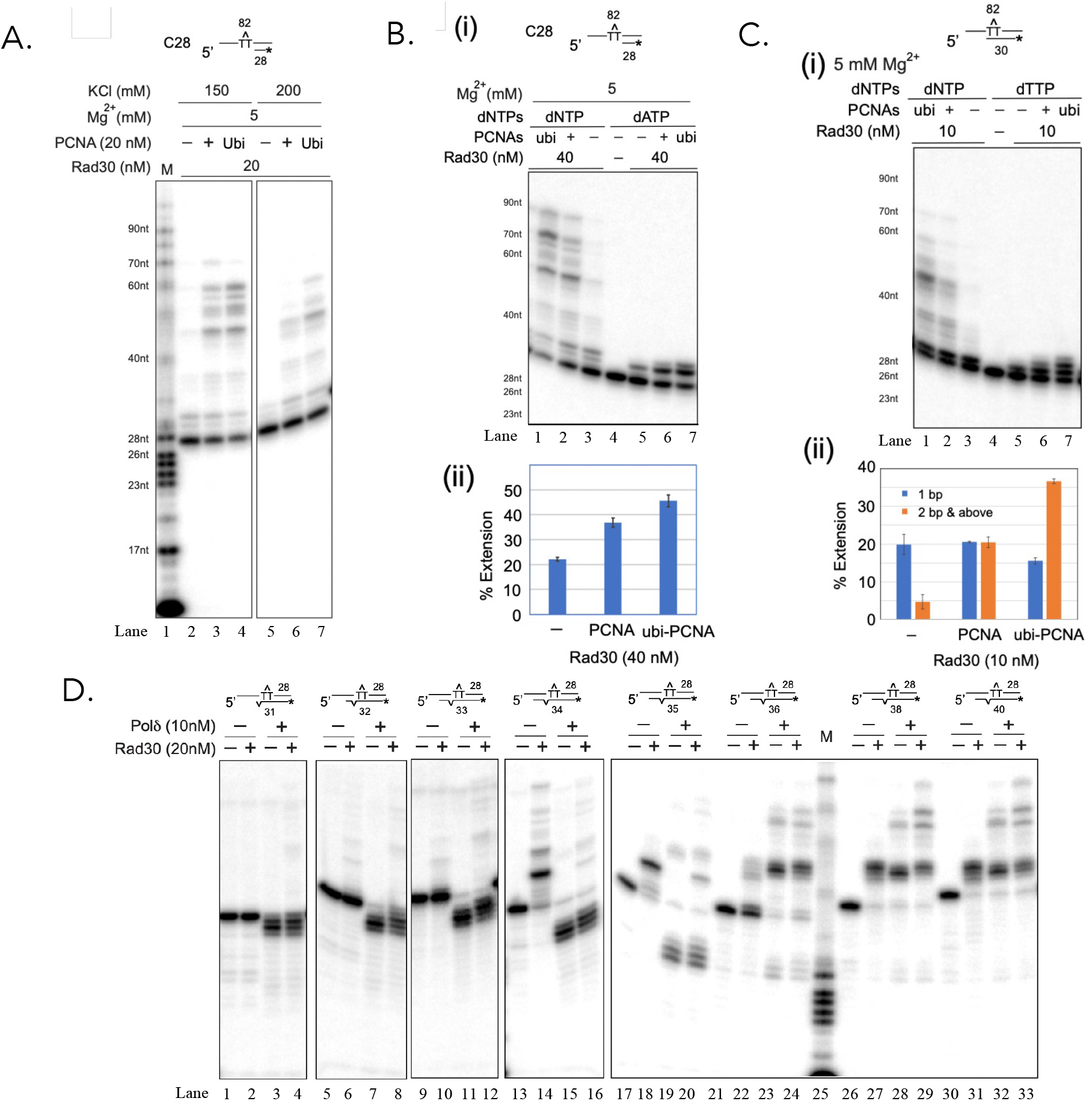
2-bp post lesion synthesis by Rad30 is inefficient, stimulated by PCNA and ubi-PCNA, and subjected to mismatch proofreading by Pol δ. **A**. Effects of PCNA and ubi-PCNA were tested in Rad30 (20 nM)-catalyzed T-T dimer bypass on the C28 substrate under 150 mM and 200 mM KCl, with RFC and RPA added to facilitate the loading of PCNA and ubi-PCNA. **B**. Similar assays were performed as in **A**, except 40 nM Rad30 was used, and reactions with only dATP were included for quantitative analysis of the lesion bypass. The results are presented in (i) and quantified in (ii). **C**. Stimulation of Rad30 (10 nM) post lesion synthesis by PCNA and ubi-PCNA were examined using the C30 substrate. Reactions with only dTTP were included to trap the 2-bp product for quantitative analysis. The results are presented in (i) and quantified in (ii). **D**. Pol δ and Rad30 were tested on a series of substrates harboring a T-T dimer on the template strand and an A/C mismatch immediately downstream of the T-T dimer. The distance between the priming site and the lesion site was varied to examine the counteraction between Rad30/Pol δ polymerization and Pol δ exonuclease digestion.

Given Rad30’s inefficient T-T dimer bypass, we investigated its activity in post-lesion synthesis using substrates representing priming positions before (C28), immediately after (C30), and two base pairs beyond (C32) the T-T dimer. Rad30 activity on the C32 T-T dimer substrate was comparable to that on the non-damaged substrate (**Fig. S9**). Notably, Rad30 activity on the C30 substrate was 2-3 times higher than on the C28 substrate, but 2-5 times lower than on the C32 substrate, suggesting reduced efficiency in synthesizing two base pairs post-lesion (**Fig. S9**). To quantitatively assess Rad30’s post-lesion synthesis, we used dTTP to trap the 2-bp product on the C30 substrate. PCNA had minimal impact on the synthesis of the first base pair but significantly enhanced the synthesis of the second and third base pairs, increasing nearly four-fold from around 5% to 20%. Ubiquitinated PCNA further boosted this rate nearly twofold, reaching over 35% (**Fig. 3C**). However, the presence of ubiquitinated PCNA also resulted in a significant amount of 3-bp products, suggesting a G/T mismatch since only dTTP was available (**Fig. 3C**). These results indicate that PCNA ubiquitination enhances Rad30’s processivity during post-lesion synthesis, although it may compromise fidelity by promoting mismatch incorporation, especially under dNTP-starving conditions.

The 3’ to 5’ exonuclease activity of Pol δ has been previously reported to efficiently remove 2-bp post-lesion synthesis products at a T-T dimer site (*32*). In our study, we observed similar results, where Pol δ exonuclease completely removed the lesion bypass products up to the 2-bp post-lesion position. Polymerization products began to appear on the primer/template substrate with a priming site 2-bp post-lesion (C32, TT+2), and Pol δ fully transitioned to its polymerase mode on the substrate with a priming site 4-bp post-lesion (C34, TT+4) (**Fig. S10**). Interestingly, when an A-C mismatch was introduced immediately after T-T dimer bypass at the TT+1 site, Rad30 exhibited reduced polymerase activity (*33*) until reaching the TT+4 site, which is 3-bp beyond the mismatch (**Fig. 3D**). In contrast, Pol δ demonstrated robust proofreading activity up to the TT+6 site, where Pol δ-catalyzed extension dominated the reaction. This indicates that Pol δ can effectively proofread mismatches up to 4-bp beyond the TT site. These findings align with genetic data, suggesting that the synchronized reduction in Rad30’s polymerase activity, coupled with increased Pol δ proofreading upon post-lesion mismatch incorporation, likely facilitates Pol δ’s proofreading for Rad30 during translesion DNA synthesis.

### Pol δ proofreading nuclease suppresses TLS at both the leading and the lagging strands

Pol δ is known as the lagging strand polymerase, but it also associates with leading strand DNA lesions when replication fork progression is perturbed (*27, 34*). This raises the question whether Pol δ exonuclease activity could inhibit TLS on both leading and lagging strands. To investigate this, we performed whole-genome sequencing (WGS) of pooled colonies from *rad14*Δ and *rad14*Δ *pol3-5DV* cells with or without UV treatment and analyze the positions of spontaneous and UV-induced mutations relative to replication origins. Because UV light induces pyrimidine dimers (Py-Py), such as TT, TC, and CC dimers, we classified single base substitution mutations into three categories: (1) mutations attributable to a single particular Py-Py (SPY), (2) mutations un-attributable to a particular Py-Py (UPY) due to the mutation located between two tandem Py-Pys and (3) mutations not associated with pyrimidine dimers, termed non-Py-Py mutations (NPY) (see **Fig. 4A** and Materials and Methods). UV treatment increased the frequency of both Py-Py-related mutations (SPY and UPY) and NPY (**Table S3**). We observed an average of 0.31 mutations per million base pairs (Mb) per colony (∼50 generations from an initial mother cell) in *rad14*Δ cells and 0.38 mutations per Mb in *rad14*Δ *pol3-5DV* cells without UV exposure, which are 5-10 fold higher than previously reported (*35*), likely reflecting the NER defect in these strains. After UV treatment, the mutation rates increased to 6.74 (21.7-fold) and 10.90 (28.7-fold) per Mb, respectively (see **Fig. 4B**).

**Fig. 4.**
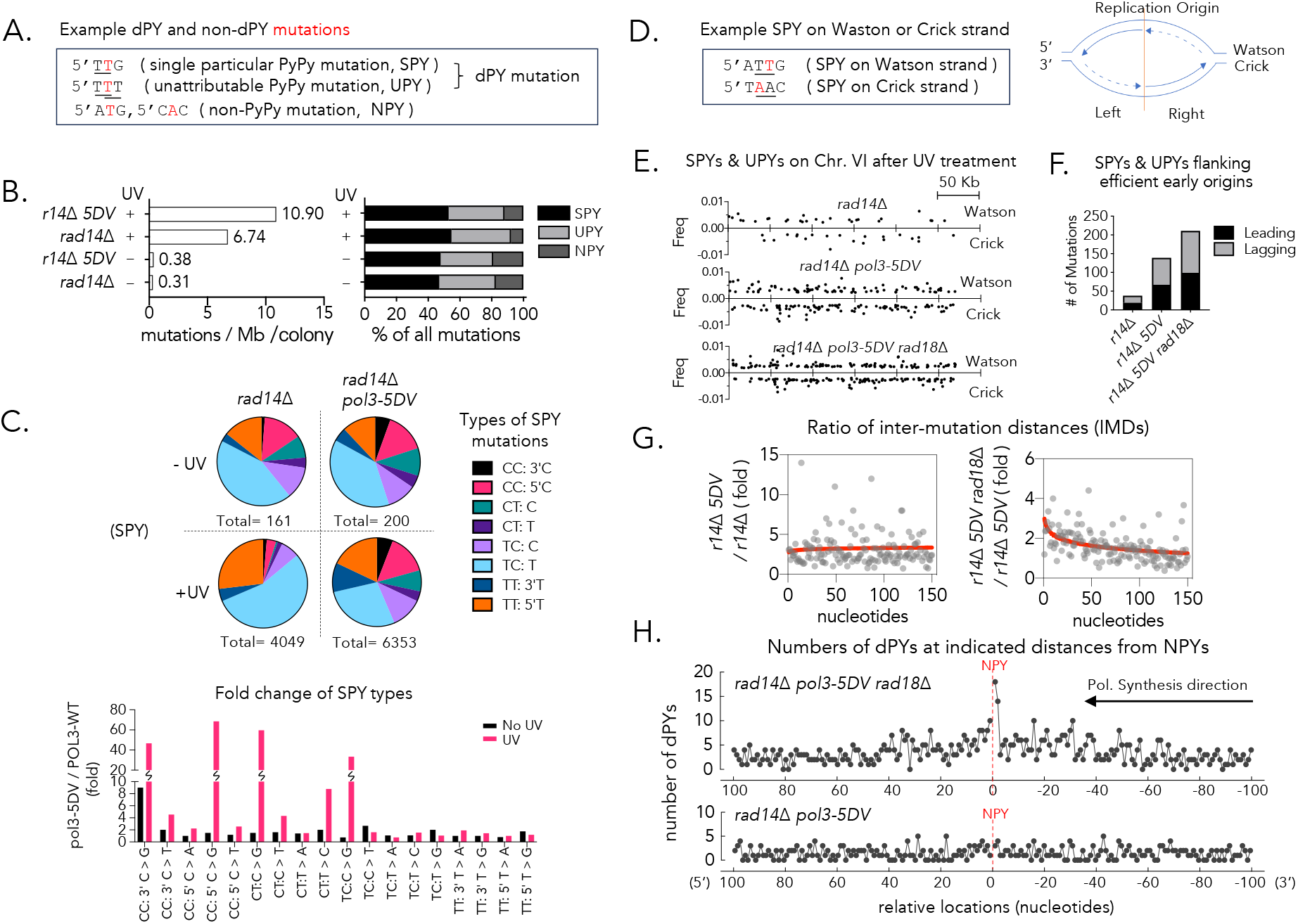
Whole genome sequencing supports that Pol δ exonuclease suppresses UV induced mutagenesis at both the leading and the lagging strands and PCNA-ubiquitylation promotes TLS extension. 100 yeast clones/survivors with or without 3 J/m^2^ UVC treatment were pooled together, and genomic DNA were isolated and subjected to whole genome sequencing at ∼2000 × coverage. **A**. Mutations can be categorized as single particular PyPy mutations (SPY), unattributable PyPy mutations (UPY), or non-PyPy mutations (NPY) based on the nucleotides adjacent to the mutations. **B**. Average number of mutations per Mb per clone in *rad14*Δ or *rad14*Δ *pol3-5DV* cells before and after UV treatment by different categories; **C**. The fold change of different types of SPY in *pol3-5DV* and Pol3 wild-type (WT) cells with and without UV treatment; **D**. Example of SPY on Watson or Crick Strand and a cartoon depicting leading and lagging strand flanking replication origin; **E**. Overview of distribution of SPYs and UPYs minus TC:T>A and TT:5’T>A mutations on the Watson and Crick strand in UV treated *rad14*Δ, *rad14*Δ *pol3-5DV* and *rad14*Δ *pol3-5DV rad18*Δ cells on chr.VI.; **F**. Increased SPY and UPY (minus TC:T>A and TT:5’T>A mutations) flanking efficient early replication origins on both the leading and the lagging strand in *pol3-5DV* cells ; **G**. The ratio of IMDs (Inter-mutation distances) at the range of 1 to 150 nucleotides between mutants as indicated. A semi-log standard curve (shown in Red) was interpolated in each graph; **H**. The numbers of dPY-NPY pairs in their relative locations (less than 100 nucleotide upstream or downstream of NPY) found in *rad14*Δ *pol3-5DV* and *rad14*Δ *pol3-5DV rad18*Δ cells.

We focused on SPY for detailed mutation profile analysis because we can assign each SPY to a specific pyrimidine dimer. We found that many types of mutations were significantly increased after UV treatment in *pol3-5DV* cells compared to *POL3 WT* cells (from ∼2 to 69-fold) (see **Fig. 4C, Table S3**), consistent with the anti-TLS role of Pol δ nuclease. Importantly, *pol3-5DV* cells exhibited a higher number of mutations globally and at both the leading and lagging strands within 5 kb regions surrounding the efficient early replication origins after UV treatment (see **Fig. 4D-F, Data S1**). These findings support the idea that Pol δ inhibits TLS on both leading and lagging strands.

### Evidence that PCNA ubiquitylation promotes TLS extension *in vivo*

Due to the low fidelity of TLS polymerases, collateral mutations can occur flanking the primary UV lesions (*36, 37*). Some of the NPY mutations could thus be induced by single TLS events. To test this possibility, we calculated the inter-mutation distance (IMD) between closely spaced mutations and compared them between *rad18* and/or *pol3-5DV* mutants. *pol3-5DV* mutation does not impact on the length of IMD within one to 150 nucleotide range (**Fig. 4G**). Interestingly, *rad18*Δ strongly increases the events with short IMD (**Fig. 4G**). The results suggest that Rad18 likely promotes TLS extension. We further analyzed the IMD between dPYs and NPYs which might be derived from pre- or post-lesion bypass of the nearby pyrimidine dimers after UV treatment (see **Data S2**). Interestingly, the dPY is strongly enriched at the first and second nucleotides immediately 3’ to NPY in *rad14*Δ *pol3-5DV rad18*Δ cells (5.0- and 3.9-fold increase respectively compared to the ±100 bp window average) (see **Fig. 4H**). The results suggest that dPY and closely spaced NPY can be induced by single TLS event by TLS polymerase(s) and the increased mutation frequency two nucleotides downstream of dPYs in *rad18* mutant can be traced to the absence of PCNA ubiquitylation and the frequent stalling of TLS polymerase(s) at +1 position, supporting our biochemical observations.

## Discussion

Collectively, these results suggest that Pol δ safeguards TLS by facilitating error-free recombination for template switching and lesion bypass. Mechanistically, Rad30’s low processivity and the frequent turnover at UV lesions allows Pol δ exonuclease activity to counteract Rad30 polymerization and ensure the fidelity of nascent TLS products. Our chromatin association results suggest that Rad18 and ubiquitinated PCNA are largely dispensable for Rad30 recruitment. Instead, ubiquitinated PCNA uniquely stimulates Rad30 to synthesize the second nucleotide downstream of the T-T dimer. We propose that PCNA and ubiquitinated PCNA enhance Rad30’s processivity, promoting both lesion bypass and subsequent 2-nucleotide post-lesion synthesis, thereby enabling Rad30 to overcome Pol δ’s exonuclease-mediated removal. It remains a question what is the true signal for recruitment of Rad30 to UV lesions. Additionally, the interaction between PCNA and TLS polymerases might help TLS polymerases outcompete replicative polymerases. Consistent with this model, biochemical reconstitution suggests that Pol δ can physically inhibit TLS by occluding the lesion-blocked 3’ end (**Fig. S11**) as reported previously (*27*). We propose that PCNA ubiquitination aids TLS polymerases in accessing blocked 3’ ends for DNA synthesis.

We found that *pol3-5DV* mutation leads to residual UV induced mutations in *rad14*Δ *tls*Δ cells (see **Fig.1G**). This prompted us to suggest that exonuclease defective Pol δ might itself acquire TLS ability to bypass DNA lesions after UVC treatment as is implicated previously (*38, 39*). Alternatively, pol3-5DV might act as an efficient “extender” after the nucleotide insertion actions by the low processive TLS polymerase Rad30. Rev3 has also been implicated as the processive extender at the AP sites (*1, 40*). We propose that multiple turnovers between DNA polymerases likely take place until TLS polymerase extends sufficiently to avoid 3’ to 5’ exonuclease activity of Pol δ, and they are stimulated by PCNA and its ubiquitylation.

Short IMDs and increased mutation occurrence at 1-2 nucleotide downstream of UV lesions in *rad14*Δ *rad18*Δ *pol3-5DV* cells also support the idea that the PCNA ubiquitylation promotes TLS extension. We can envision that frequent mutations made during TLS extension may trigger the dissociation of TLS polymerase and the switch to Pol δ that subsequently proofreads mismatches to avoid excessive mutagenesis during TLS extension.

It is puzzling that the leading-strand replicative Pol ε does not invoke TLS suppression. Reconstitution of yeast replisome suggests that besides lagging strand synthesis, Pol δ also plays a role in establishing leading-strand synthesis before DNA polymerase ε engagement (*41, 42*). This raises a possibility if cells might employ Pol δ preferentially following TLS even during the leading strand lesion bypass and engages Pol ε only after Pol δ polymerase to successfully switch on from TLS pols. Alternatively, most TLS might take place at post-replicative gaps, where Pol δ was shown to function primarily for gap fill-in synthesis (*43, 44*).

The expression of Rad18 and TLS polymerases is frequently elevated in many cancers and during targeted therapy (*45, 46*). Mutations in Pol δ region at or near proofreading nuclease domain are also found in many cancers including breast, colorectal and endometrial cancers (*47-49*). TLS is crucial for the viability of cancer cells, particularly those deficient in recombination (*50, 51*). Our study highlights a novel role for DNA proofreading nuclease in limiting low-fidelity lesion bypass catalyzed by TLS polymerases, thereby reducing mutagenesis, and promoting recombinational repair during DNA damage tolerance. The elevated Rad18 expression might counteract anti-TLS and pro-recombinational actions by Pol δ proofreading nuclease and could shift balance toward TLS. It will be important to determine whether similar mechanisms operate in human cells and whether elevated TLS affect cancer evolution and therapy resistance in cancers deficient in Pol δ nuclease activity, potentially offering new opportunities for improved cancer treatments.

## Acknowledgments

We are grateful to Dmytri Gordenin, Thomas Kunkel ,Patrick Sung,Michael Fasullo and Xiaolan Zhao for providing plasmids and yeast strains, and Dmytri Gordenin for critical reading of the manuscript.

## Fundings

This work was supported by William and Ella Owens Medical Research Foundation, and NIH research grant GM141631 to S.E.L., National Key Research and Development Program 2021YFA0909300 & High level of special funds G03050K003 to H.R., American Cancer Society Research Scholar Award RSG-21-013-01-DMC and NIH R35 research grant GM152207 to H. N., and NIH R35 research grant GM 127029 to J.E.H. .D.N.G. was supported by NIGMS Genetics Training Grant T32GM007122, and by the National Science Foundation Graduate Research Fellowship Program under grant 1744555.

## Author contributions

Conceptualization: J.C., S.E.L. and H.N.; Data collection and investigation: J.C., Q.W., J. H., and D.N.G.; Methodology: J.C., Q.W., J.H., Z.Z., J.M.C. and D.N.G.; Funding acquisition: S.E.L., H.N., H.R., X.H. and J.E.H.; Supervision: E.S., S.E.L. and H.N.; Writing – original draft: J.C. and Q.W.; Writing – review and editing: J.C., S.E.L., H.N. and J.E.H.

## Supplementary Materials

Materials and Methods Figs. S1 to S12

Tables S1 and S4 Data S1 to S2

References (*52*–*63*)

## Notes

### Competing Interest Statement

The authors have declared no competing interest.

