## Supplemental Materials for "Yeast DNA Polymerase δ Proofreading Nuclease Suppresses Translesion Synthesis and Facilitates Recombination in DNA Lesion Bypass"

**The PDF file includes:**

Materials and Methods
Supplementary Text
Figs. S1 to S12
Tables S1 to S4
References 52-63

**Other Supplementary Materials for this manuscript include the following:**

Data S1 to S2

### Materials and Methods

#### Yeast strains

Yeast strains were constructed either by standard one-step gene replacement using PCR products to modify strains YKJM1(*52*), YB163(*53*) and TGI354(*54*), or by sporulation to generate various genotypes in W303(*55*) isogenic strains. Point mutations in *POL2*, *POL3* and *RAD30* at their genome loci were generated by two-step gene replacement by a *URA3* plasmid integration and then FOA selection for $Ura^-$ pop-outs. Mutants were identified by PCR with an oligo that anneals to either wild-type or the mutant sequence at its 3' terminus. The genotype of strains used in this study are listed in Table S1.

#### Cells cultures and treatment

#### Cell survival assay

Single colonies of yeast cells grown overnight were subjected to five-fold serial dilution in 96-well plates and spotted onto YEPD plates with or without a DNA damaging agent, as indicated. For UV treatment, cells plated on YEPD plates were treated with 254 nm UVC with a hand-held UV irradiator fixed at certain distance from plate at a dose rate of 0.4-1 $J/m^2/sec$ for 10~60 $J/m^2$ or at 0.1-0.2 $J/m^2/sec$ for 2-5 $J/m^2$ UV. Cells were then incubated in dark for 2-5 days at 30°C before taking pictures.

#### UV-induced unequal sister chromatid recombination (uSCR) assay

Single colonies of yeast cells bearing a uSCR cassette(*53*) were grown in YEPD overnight to stationary phase and the spontaneous frequency of uSCR was determined using each of the cell cultures by plating on synthetic media (SM) lacking histidine. Appropriate dilutions of the cell cultures were plated on YEPD plates in parallel to determine the total number of cells plated on SM-histidine. 140 µl of overnight yeast cultures were then inoculated in 14 ml of fresh YPED and grown for 6 h to establish logarithmic growth. Cells were then harvested, washed once with 10 ml double-distilled $H_2O$ ($ddH_2O$, resuspended in 14 ml of $ddH_2O$ and transferred to a 9 cm petri dish. Cells were then irradiated for 50 $J/m^2$ UV at 2 $J/m^2/s$ on top of a horizontal shaker with slow rotating. Cells were then collected from the petri dish and plated on SM-histidine as well as YEPD plates with appropriated dilutions. The UV-induced uSCR frequency was calculated by subtracting the spontaneous uSCR in each corresponding cell culture.

#### Spontaneous and UV induced *CAN1* mutagenesis

Single colonies of yeast cells were grown in YEPD overnight to stationary phase and the spontaneous frequency of $can^r$ clones was determined by plating on canavanine-containing SD plates and on YEPD plates with appropriated dilutions.

To determine UV-induced mutation frequencies, *rad14Δ* derivative cells (Fig. 1G) were arrested in G1/S boundary using alpha factor (10μg/ml), washed once with $ddH_2O$, suspended in 14 ml of water in a 9 cm petri dish and then irradiated with 3 $J/m^2$ UVC at dose rate of 0.2 $J/m^2/s$. Cells were then grown in YEPD for 4-5 hours before plating on SD + canavanine and YEPD plates with appropriate dilutions. For *RAD14+* cells (Fig. 2G), 1.5 ml of stationary phase cultures were washed with $ddH_2O$, suspended in 14 ml of water in a 9-cm petri dish, irradiated with 30 $J/m^2$ UV at dose rate of 2 $J/m^2$/sec and then grown in 10 ml YEPD overnight before plating on SD + canavanine and YEPD plates with appropriate dilutions. The UV-induced $can^r$ frequency were calculated by subtracting the spontaneous frequency in each corresponding cell culture.

### Whole-genome deep sequencing and analysis of UV-induced mutations

**Sample preparation:** For whole genome deep sequencing, appropriate numbers (~ 200 or 2000) of BY4741 *rad14Δ* and *rad14Δ pol3-5DV* cells were plated on YEPD plates and treated with or without 3 $J/m^2$ UV at a dose rate of 0.15 $J/m^2$/sec. 100 surviving colonies from each genotype were pooled and genomic DNA were then isolated as previously described(*56*). These samples were subjected to high-throughput whole genome deep sequencing to about 2000 × coverage of yeast genome size (~30 Gbp) using a PCR-free DNBseq platform at Beijing Genome institute (BGI). We assume that UV-induced DNA lesions are randomly distributed and not enriched in certain regions of the genome. Due to the relative low UV dosage, the majority of UV lesions are unique in individual colonies. Therefore, a mutation generated in the mother cell of one colony after TLS should be present in about 0.5 % of mutant reads among wild-type reads because 1 daughter cell contains a mutation while the other daughter cell contains wild-type allele) and because we mixed 100 individual colonies. There are i.e. about 10 mutant reads among about 2000 wild-type reads on average.

**Variant calling** The quality control of raw sequencing data was performed with FastQC (https://www.bioinformatics.babraham.ac.uk/projects/fastqc/). Adapters and low quality reads were filtered using Trim Galore (https://www.bioinformatics.babraham.ac.uk/projects/trim_galore/). Trimmed reads were aligned to the *Saccharomyces cerevisiae* reference genome (R64-1-1) using BWA-mem with default parameters(*57*). To ensure data quality and retain only high confidence reads, only paired-end aligned reads were kept, using Samtools(*58*). To ensure mate-pair information was consistent, FixMateInformation was carried out with GATK v4.4.0.0. (doi: https://doi.org/10.1101/201178)(*59*). Variants were identified using GATK's HaplotypeCaller with following parameters according to GATK's : --pcr-indel-model NONE, --stand-call-conf 50, --sample-ploidy 200. SNPs with allele count (AC)=1 were retained and further quality control was performed according to GATK's hard-filtering manual.

**Data analysis:** Based on reference genome sequence (BY4741, Saccharomyces genome database), individual mutations were assigned to different mutation categories using R script based on following rules: if a mutation is on a pyrimidine (or purine) and its 5' or 3' nucleotide is also a pyrimidine (or purine), then this mutation is defined as a single particular Py-Py mutation (SPY). If both 5' and 3' nucleotides of a pyrimidine (or purine) mutation are pyrimidines (or purines), then this mutation is defined as unattributable Py-Py mutation (UPY). All remaining mutations were assigned as non-Py-Py mutations (NPY). For Fig.4E & 4F, SPY and UPY mutations 5 kb flanking efficient early origins were sorted out and allocated the leading (Left top + Right bottom) and lagging strands (Left bottom + Right top) (table S4). The TC: T>A (an abbreviation of the 5'TC dimer with its T mutated to A) and TT:5'T>A mutations were excluded from the analysis in Fig. 4E & 4F due to their apparent lack of proofreading by Pol δ, judging by their frequency are decreased in *pol3-5DV* cells after UV treatment (see Table S3). The inter-mutation distances (IMDs) were calculated between each neighboring mutation and the IMDs ranging from 1-150 nt were sorted out. The ratio of IMDs between different samples were then calculated and plotted as a graph in Fig 4G. A semi-log standard curve was interpolated in each graph.

dPYs flanking all NPYs in samples with ⩽ 150nt distances were identified and the relative locations of dPY (3' or 5' to NPY) as well as the their strand location (top or bottom) were used to determine whether the dPY is at upstream or downstream of NPY with regards to the moving direction of TLS polymerase (s).

**Fractionation of formaldehyde-crosslinked chromatin**

Chromatin fractionation assay was based on a previous protocol(*60*) with some key changes. For UV treatment, log-phase yeast cells (1-2 ×$10^7$ cells/ml) were irradiated with 100 J/$m^2$ UV in dd$H_2O$ and cultured in YEPD for 1 h. 10 ml of UV-irradiated or non-irradiated cells were treated with 1% formaldehyde for 15 min and then neutralized with 0.25 M glycine for 5 min on a shaker. Cells were then washed with 10 ml of Tris Buffered Saline (20mM Tris pH8.0, 150mM NaCl) and frozen at -80°C. Cells were suspended in 500 µl of pre-spheroplast buffer (1M sorbitol, 50mM EDTA, 2% β-mercaptoethnol, 2 mM PMSF) and incubated at room temperature for 5 min before spun down. Cell pellets were then suspended in 500 µl spheroplast buffer (1 M sorbitol, 50 mM EDTA) containing 0.75 mg/ml Zymolyase 100T, 2% β-mercaptoethnol, 2 mM PMSF and 1.5 × proteinase inhibitor cocktail and incubated at 30°C for 20 min. The spheroplasts were then spun down and washed once with spheroplast buffer. 100 µl of ice-cold pre-lysis buffer (100 mM Tris pH8.0, 50 mM EDTA, 100 mM NaCl, 1× proteinase inhibitor cocktail, 2 mM PMSF) were gently added on top of the spheroplast pellets and spun with rotating the tubes 180°in the centrifuge at 6000 rpm at 4°C for 2 min to remove residual sorbitol. The spheroplasts were then resuspended in 200 µl of lysis buffer (100 mM Tris pH8.0, 50 mM EDTA, 100 mM NaCl, 1% Triton X-100, 1× proteinase inhibitor cocktail, 2 mM PMSF). 30 µl of 10% N-lauroysarcosyl were then added and pipetted a few times and then treated with 5 µl of RNaseA (10 mg/ml) at room temperature for 10 min. 100 µl of ice-cold 30% sucrose (containing 1× proteinase inhibitor cocktail and 2mM PMSF) were then slowly added to the bottom of tubes. The tubes were then spun down at 13200 rpm for 10 min at 4°C. The chromatin pellet was washed once with 100 µl lysis buffer by rotating the tubes 180° in the centrifuge and then resuspended in 1×SDS Laemmli buffer and boiled at 100°C for 10 minutes.

**Expression and purification of Flag-Rad30 from *S. cerevisiae*.**

To express Flag-Rad30 in yeast, the *RAD30* coding sequence was cloned between the NotI and SpeI restriction sites of the pESC-URA vector (Agilent), placing the gene under the control of the

*GAL10* promoter and fusing a FLAG tag to the N-terminus of Rad30. The protease-deficient yeast strain 334 (*MATα pep4-3 prb1-1122 ura3-52 leu2-3,112 reg1-501 gal1*) was used for expression. For induction of Rad30 expression, an overnight culture of yeast was diluted 1:50 into 12 liters of Ura- medium containing 2% glucose. The culture was incubated at 30°C until the optical density at 600 nm reached 0.8. At this point, galactose was added to a final concentration of 2% to induce expression. After 12 h of further incubation, the cells were harvested by centrifugation, and the resulting pellet (approximately 50 g) was stored at -80°C. For protein extraction, the frozen yeast pellet was ground with dry ice, and 50 ml of K buffer (20 mM $KH_2PO4$, pH 7.4, 10% glycerol, 0.5 mM EDTA, 0.1% NP-40, 1 mM β-mercaptoethanol) containing protease inhibitors (5 μg/ml each of aprotinin, chymostatin, leupeptin, and pepstatin A, and 1 mM phenylmethylsulfonyl fluoride) and 500 mM KCl was added to the ground extract. All steps were performed at 0–4°C. The extract was clarified by centrifugation at 20,000g for 20 minutes and incubated with 0.4 ml of anti-FLAG M2 resin for 2 hours with gentle mixing. The resin was then washed sequentially with 30 ml of K buffer containing 500 mM KCl and 0.1% NP-40, followed by 10 ml of K buffer containing 500 mM KCl. Flag-Rad30 was eluted from the resin using 1.5 ml of K buffer containing 500 mM KCl and 200 μg/ml FLAG peptide for 1 hour. The eluate containing purified Rad30 was subjected to filter dialysis and concentrated using an Ultracel-30K concentrator (Amicon) to a final concentration of 1 mg/ml. The purified protein was stored in aliquots at -80°C. The overall yield of Rad30 protein was approximately 200 μg.

**Yeast DNA Pol δ, replication factor C, PCNA and replication protein A.**

The yeast DNA polymerase δ complex, including the exonuclease-deficient *pol3-01*mutant (D321A, E323A), was expressed in the protease-deficient yeast strain BJ5464 (*MATα pep4::HIS3 prb1Δ1.6R ura3-52 trp1 leu2Δ1 can1 GAL*). This complex consisted of FLAG-tagged Pol3 (pBJ1445), GST-tagged Pol31, and Pol32 (pBJ1524) (Kind gifts from Dr. Patrick Sung). Site-directed mutagenesis was used to introduce the D321A and E323A point mutations into the Pol3 gene in plasmid pBJ1445 to generate the exonuclease-deficient mutant. Similarly, the Replication Factor C (RFC) complex, consisting of Gst-Rfc1, Rfc2, Rfc3, Rfc4, and Rfc5, was expressed in the same yeast strain (BJ5464). For expression of RFC, plasmid pBJ1476 (2 μ, *GAL-PGK-GST-RFC1/RFC4/RFC5, LEU-2d*) was used for RFC1, RFC4, and RFC5 (Kind gifts from Dr. Patrick Sung), while plasmid pBJ1469 (2 μ, *GAL-PGK-RFC2/RFC3, TRP1*) was used for RFC2 and RFC3. Proliferating cell nuclear antigen (PCNA) was expressed in *E. coli* using plasmid pQE16-Pol30 (a kind gift from Dr. Patrick Sung). All above proteins were purified using previously described protocols(*61*).The single-stranded DNA-binding protein (RPA) was expressed in yeast and purified following established methods(*62*).

**Yeast Ubi-PCNA.**

The N-His-mono-ubiquitin-*PCNA* coding sequence was synthesized and cloned into the pET21a(+) vector using NdeI and XhoI restriction sites by Gene Universal (sequence available upon request). The *Rosetta2, pLysS* strain was transformed with the ubi-PCNA expression vector in *E. coli*. For protein expression, an overnight culture was diluted 1:50 into 4 liters of LB medium containing 100 mg/l ampicillin and 37 mg/l chloramphenicol. When the culture reached an optical density of A600 = 0.8, expression of Ubi-PCNA was induced by adding 0.5 mM IPTG, followed by overnight incubation at 16°C. After centrifugation, a cell pellet of approximately 10 g was obtained. For purification, the cell pellet was resuspended in 50 ml of K buffer (containing 150 mM KCl) and disrupted by sonication for 4 min, using a cycle of 2 sec of sonication followed by 4 seconds of rest. The lysate was clarified by centrifugation at 20,000g for 20 min and applied to a 5 ml Q Sepharose column (GE). The column was washed with 100 ml of K buffer containing 150 mM KCl. Ubi-PCNA was eluted using 30 ml of K buffer containing 500 mM KCl and was gently mixed with 0.4 ml of Ni-NTA resin for 2 h. The resin was then washed with 30 ml of K buffer containing 500 mM KCl, 0.1% NP-40, and 15 mM imidazole, followed by 10 ml of K buffer with 500 mM KCl and 15 mM imidazole. Finally, Ubi-PCNA was eluted with 3 ml of K buffer containing 500 mM KCl and 200 mM imidazole. The eluate containing purified Ubi-PCNA was filter-dialyzed and concentrated using an Ultracel-30K concentrator (Amicon) to a final concentration of 10 mg/ml. The protein was stored at −80°C in aliquots, with an overall yield of approximately 1 mg.

***DNA substrates.***
The oligonucleotides used to construct substrates for biochemical assays are listed in Table-S4. The Cis-Syn Thymidine Dimer-containing oligonucleotide (H2) was synthesized by Trilink Biotechnologies, while all other oligonucleotides were purchased from IDT. To generate 5′,3′ double-overhanging substrates, primers of varying lengths (H3–H10) were 5′-labeled with [$\gamma$-$^{32}$P] ATP (PerkinElmer) using T4 polynucleotide kinase (New England Biolabs) and annealed to either the undamaged template (H1) or the Thymidine Dimer-containing template (H2).

**Rad30 translesion and Pol δ proofreading assay**
Unless otherwise specified, DNA substrate (10 nM) was incubated with 10 nM RFC, 20 nM PCNA or mono-ubiquitin-PCNA, and 30 nM RPA in a 9 µl reaction buffer (20 mM Tris-HCl, pH 8.0, 2 mM $MgCl_2$, 1 mM DTT, 200 ng/ml BSA, 150 mM KCl, and 1 mM ATP) for 10 min at 30°C. Subsequently, 1 µl of Rad30 or Pol δ, at the indicated concentration, was added, and the reaction was incubated for an additional 10 min at 30°C. To allow full extension by Rad30 or Pol δ, 0.1 mM of each of the four dNTPs was added to the reaction. Alternatively, to trap a limited extension product (2–3 nucleotides in length) for quantification, only individual dNTPs were added. Reactions were stopped by adding an equal volume of loading dye containing 85% formamide and 25 mM EDTA, along with 1 µM unlabeled H11 oligonucleotide to prevent the re-annealing of radiolabeled primers to the complementary strand. The mixtures were then heated at 95°C for 5 min and fractionated on a 12% denaturing polyacrylamide gel in TBE (Tris-borate-EDTA), followed by phosphorimaging analysis.

**Rad30 heparin trapping experiment**
A heparin-trapping experiment (*63*) was performed to assess the single-turnover activity of Rad30. DNA substrate (10 nM) was incubated with 30 nM Rad30 in a 9 µl reaction buffer (20 mM Tris-HCl, pH 8.0, 2 mM $MgCl_2$, 1 mM DTT, 200 ng/ml BSA, 150 mM KCl) for 10 min at 30°C to allow Rad30 binding to substrate DNA. Following this, 1 µl of 1 mM dNTP and 1 mg/ml heparin was added to start the reaction and trap unbound Rad30. The reaction was incubated for an additional 10 min at 30°C, stopped, and the products were gel-fractionated and analyzed via phosphorimaging.

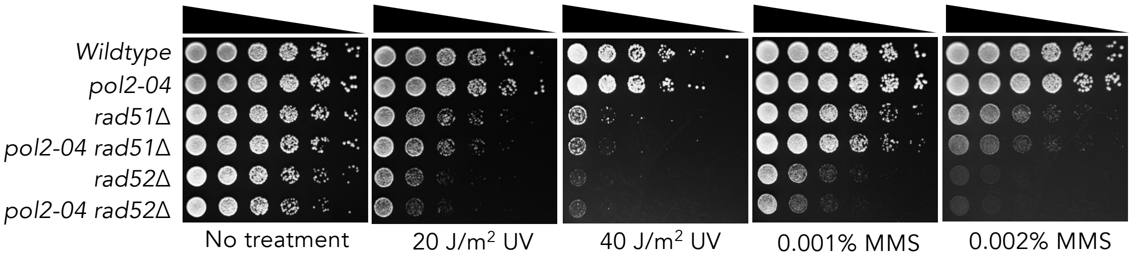

**Fig. S1. Pol ε proofreading mutant does not suppress UV or MMS sensitivity of cells deficient in homologous recombination.** Sensitivity of cells with indicated genotypes to different types of DNA damage. 5-fold serial dilutions of cells were spotted onto YEPD plates and treated with or without various UV doses or YEPD plates containing indicated concentrations of MMS. Cells were then incubated in dark for 2-5 days at 30° C before taking pictures.

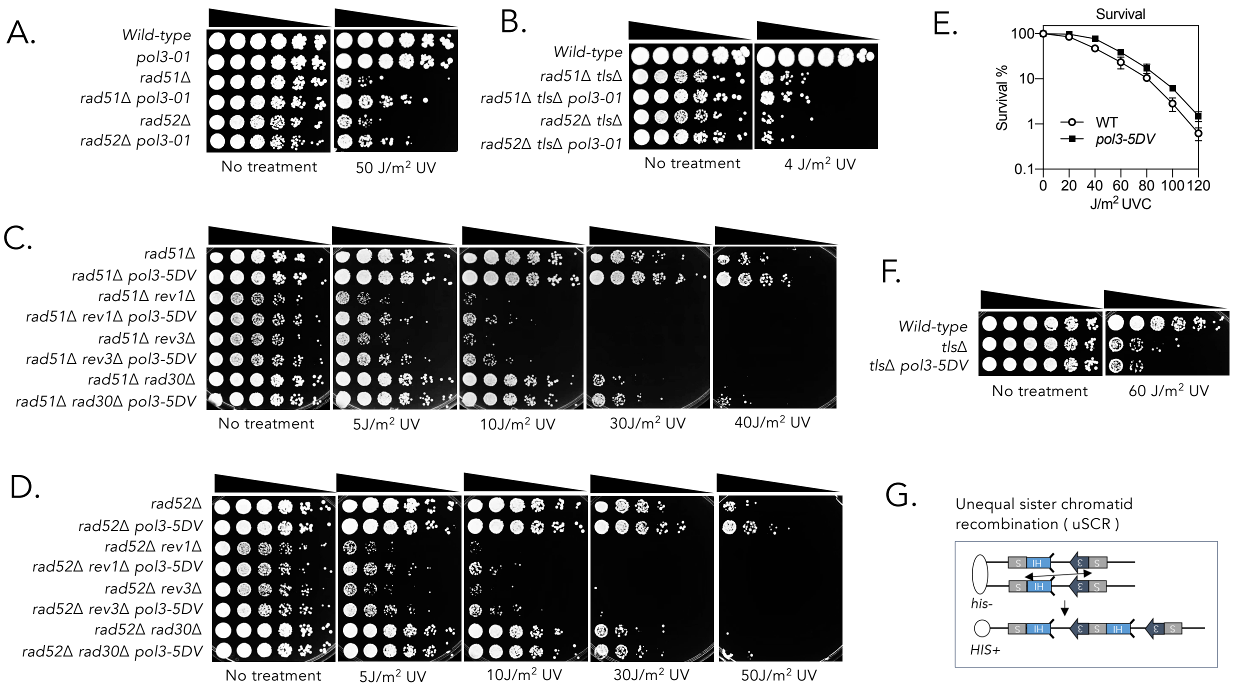

**Fig. S2. The suppression of UV sensitivity by Pol δ proofreading mutation in homologous recombination mutants depends on all TLS polymerases. A-D&F.** Sensitivity of cells with indicated genotypes upon different doses of UV irradiation. 5-fold serial dilutions of cells were spotted onto YEPD plates and treated with or without indicated amount UV; **E.** Percent survival of wild-type and *pol3-5DV* cells to different doses of UV; **G.** The schematic principle of a genetic reporter that measure the frequency of unequal sister-chromatid recombination (uSCR). The reporter contains tandem *his3* repeats, *his3*-Δ*5*′ and *his3*-Δ*3*′ *::HOcs* at the *trp1* locus. An unequal recombination between two sister chromatids within this region would lead to intact *HIS3* gene, thus cell becomes histidine autotroph.

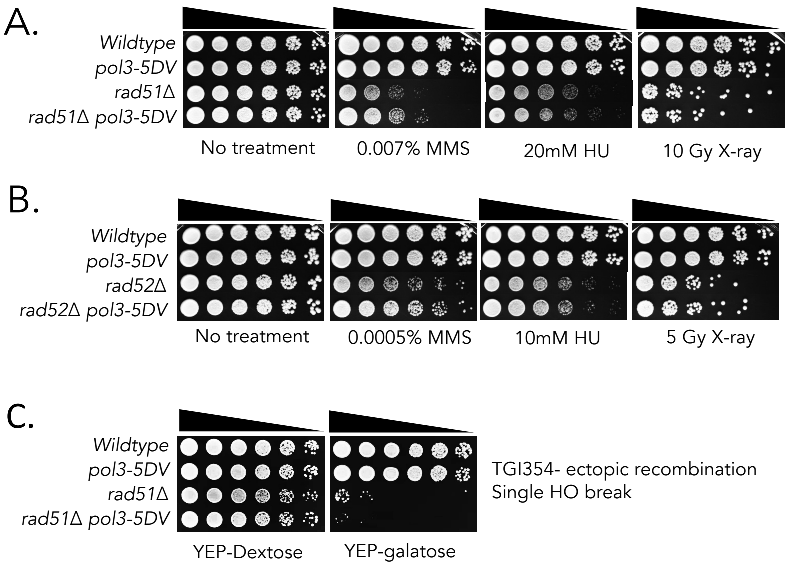

**Fig. S3. Pol δ proofreading mutation does not suppress the sensitivity to HU, X-ray or a HO-induced DSB in cells deficient in homologous recombination. A-E.** Sensitivity of cells with indicated genotypes to different DNA damage types. 5-fold serial dilutions of cells were spotted onto YEPD plates and treated with or without various amount DNA damage or YEPD plates containing indicated concentrations of DNA damage agents. Cells were then incubated in dark for 2-5 days at 30° C before taking pictures.

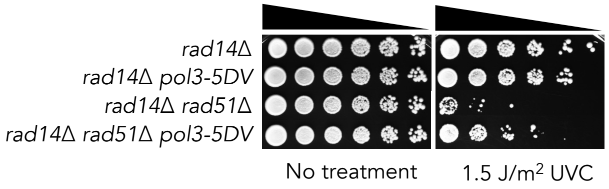

**Fig. S4. Pol δ proofreading mutation suppresses UV sensitivity of *rad51Δ* cells independently of nucleotide excision repair.** Sensitivity of cells with indicated genotypes to UV treatment. 5-fold serial dilutions of cells were spotted onto YEPD plates and treated without or with indicated amount of UV. Cells were then incubated in dark for 2-3 days at 30° C before taking pictures.

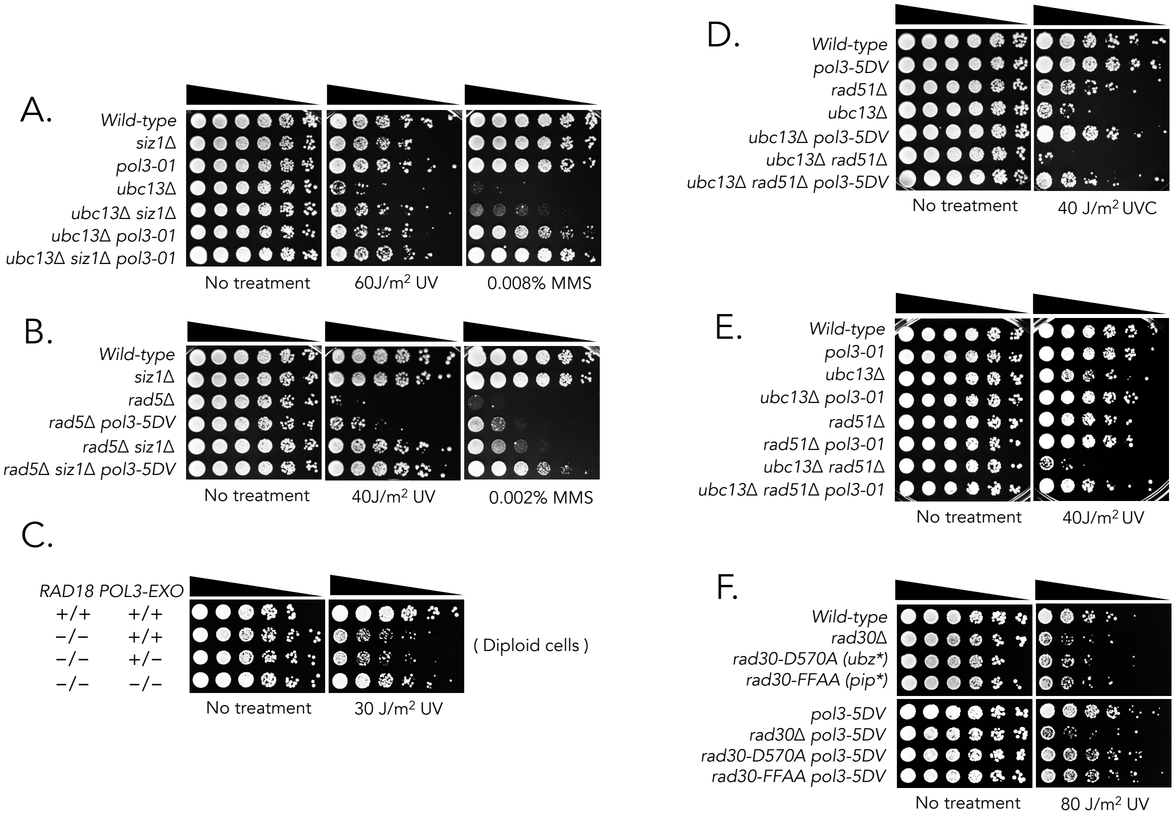

**Fig. S5. Pol δ proofreading mutation reduces UV and MMS sensitivity of cells defective in PCNA ubiquitylations by the mechanism different from *siz1Δ* and Rad51*;* A-F.** Sensitivity of cells with indicated genotypes to different types and doses of DNA damage. 5-fold serial dilutions of cells were spotted onto YEPD plates and treated with or without indicated amount of UV or MMS. Cells were then incubated in dark for 2-5 days at 30° C before taking pictures.

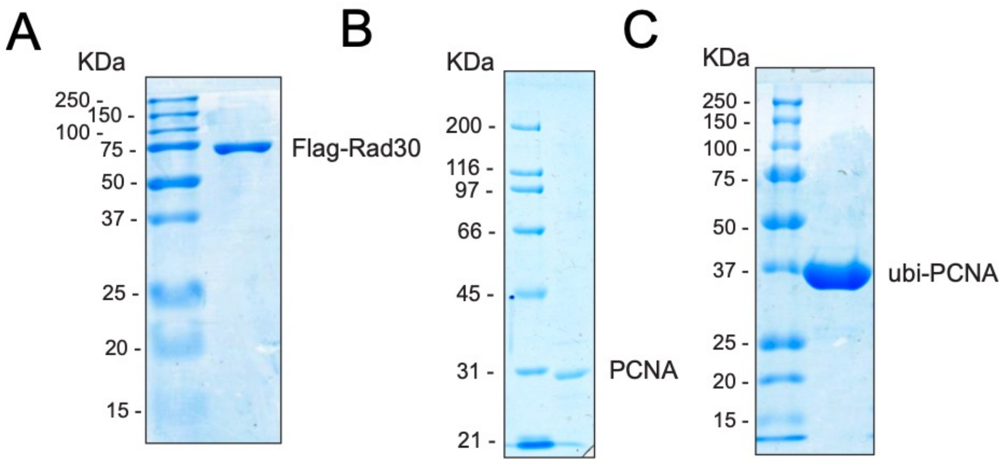

**Fig. S6. SDS-PAGE analyses of purified proteins.**

Purified Rad30 (**A**), PCNA (**B**) and ubi-PCNA (**C**) were analyzed by SDS-PAGE followed by Coomassie staining.

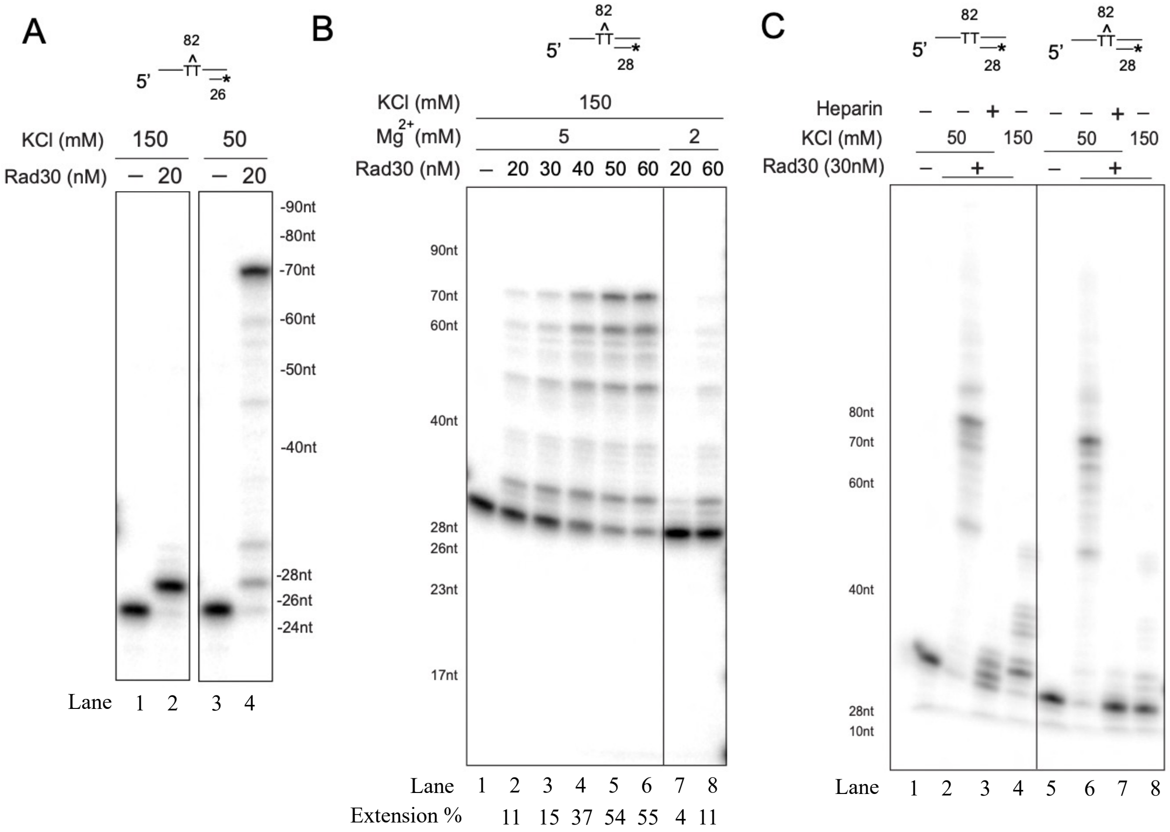

**Fig. S7. T-T dimer bypass by Rad30 requires multi-turnover. A.** Comparison of the effect of salt concentration (150 mM and 50 mM KCl) on Rad30-catalyzed T-T dimer bypass from the priming site 2 nucleotides prior to the lesion site (C26). **B.** Comparison of the effect of $Mg^{2+}$ concentration (5 mM and 2 mM) on a titration of Rad30 (20 nM to 60 nM) in the T-T dimer bypass reaction primed immediately prior to the lesion site (C28). **C.** Rad30-catalyzed primer extension was examined on both non-damaged and T-T dimer templates under varying salt conditions (50 mM and 150 mM KCl), with or without heparin treatment.

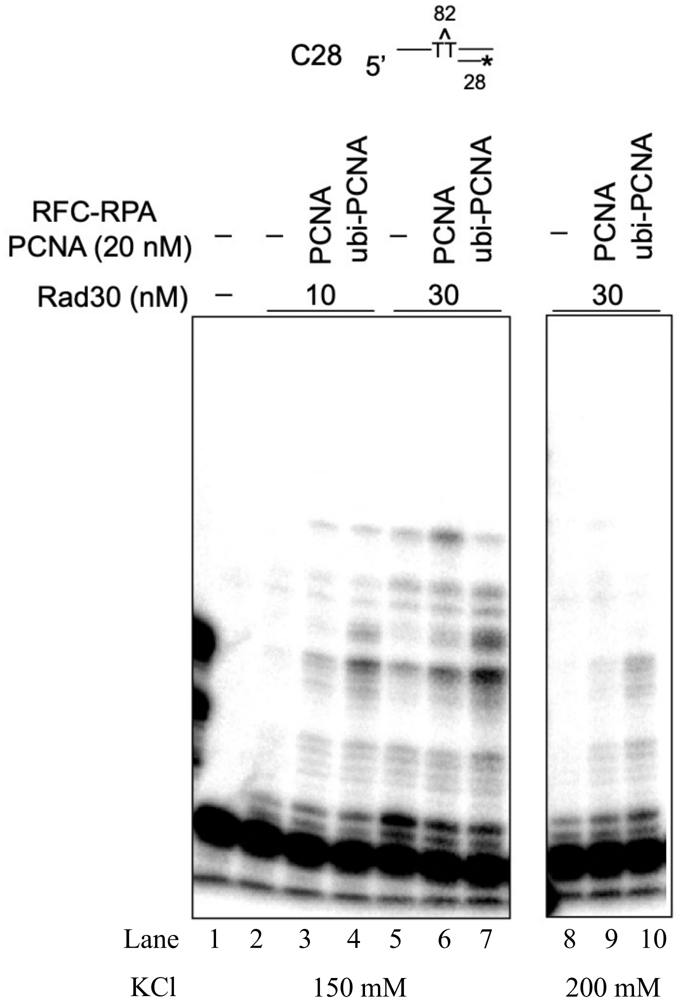

**Fig. S8. Stimulation of Rad30 catalyzed T-T dimer bypass under 2 mM $Mg^{2+}$ conditions.** T-T dimer bypass by Rad30 (10 nM and 30 nM) was tested on a 5' radiolabeled primer/template substrate (C28) under 2 mM $Mg^{2+}$ and various salt conditions, both without and with PCNA or ubi-PCNA, as well as RFC and RPA. The reactions were analyzed by denaturing polyacrylamide gel electrophoresis and phosphoimaging.

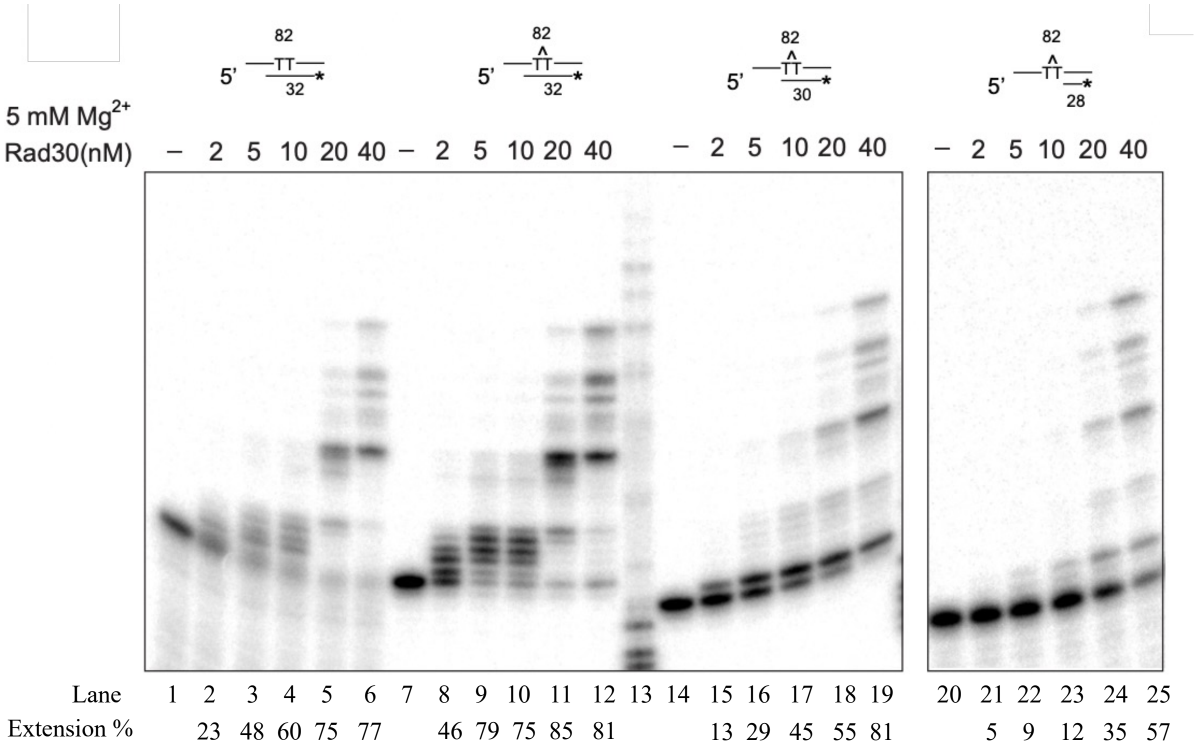

**Fig. S9.** Reduced efficiency in synthesizing two base pairs post-lesion by Rad30.

Lesion bypass and post-lesion synthesis by Rad30 were compared using Rad30 titration against primer/T-T dimer template substrates with priming sites positioned immediately before the lesion (C28), after the lesion (C30), and 2 bp past the lesion (C32). A 2-bp post-lesion substrate with a non-damaged template was also included for comparison.

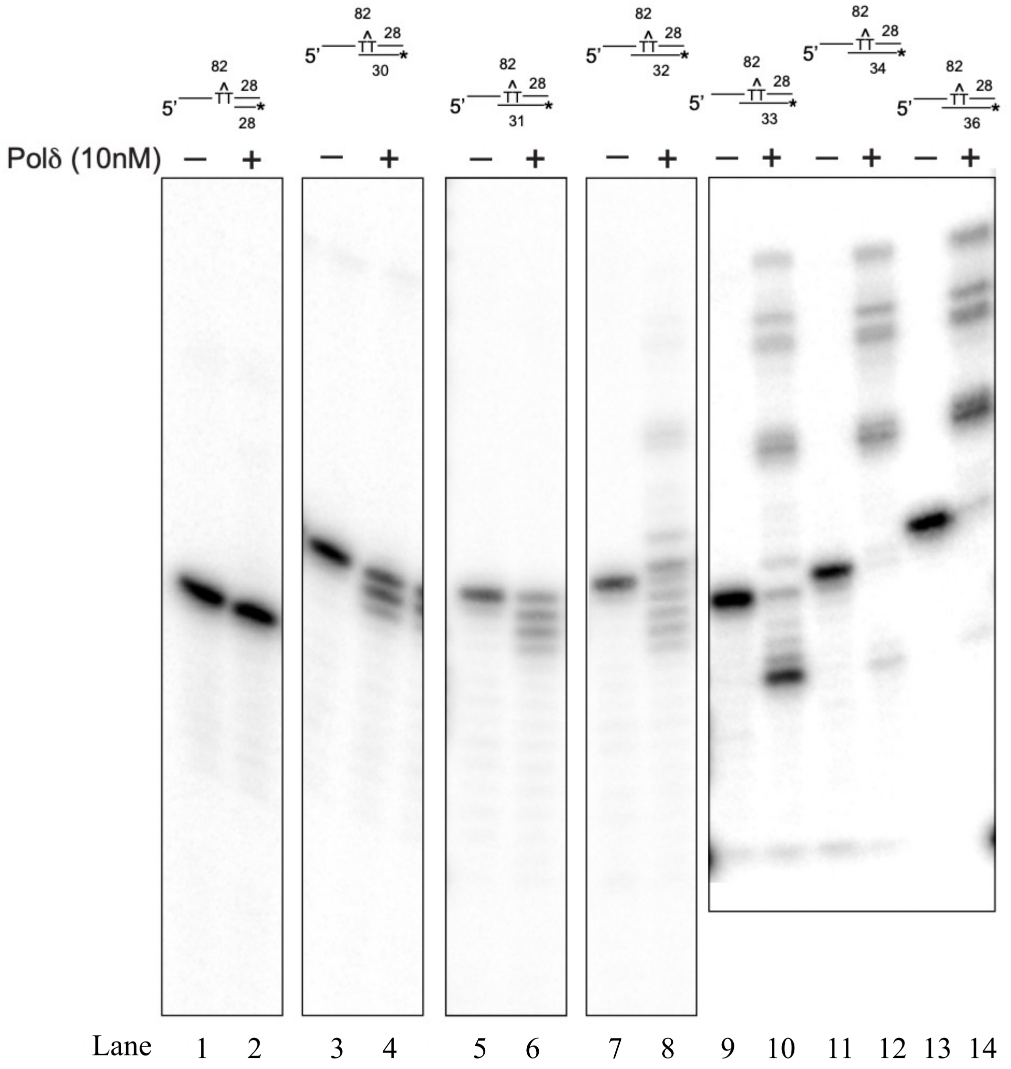

**Fig. S10. Switching of Pol δ from exonuclease digestion to polymerization depends on the distance of the priming site surpassing the T-T dimer.** Pol δ (10 nM) was reacted with 5' radiolabeled primer/template substrate with the priming site prior to the T-T dimer site, or at an indicated distance post T-T dimer bypass.

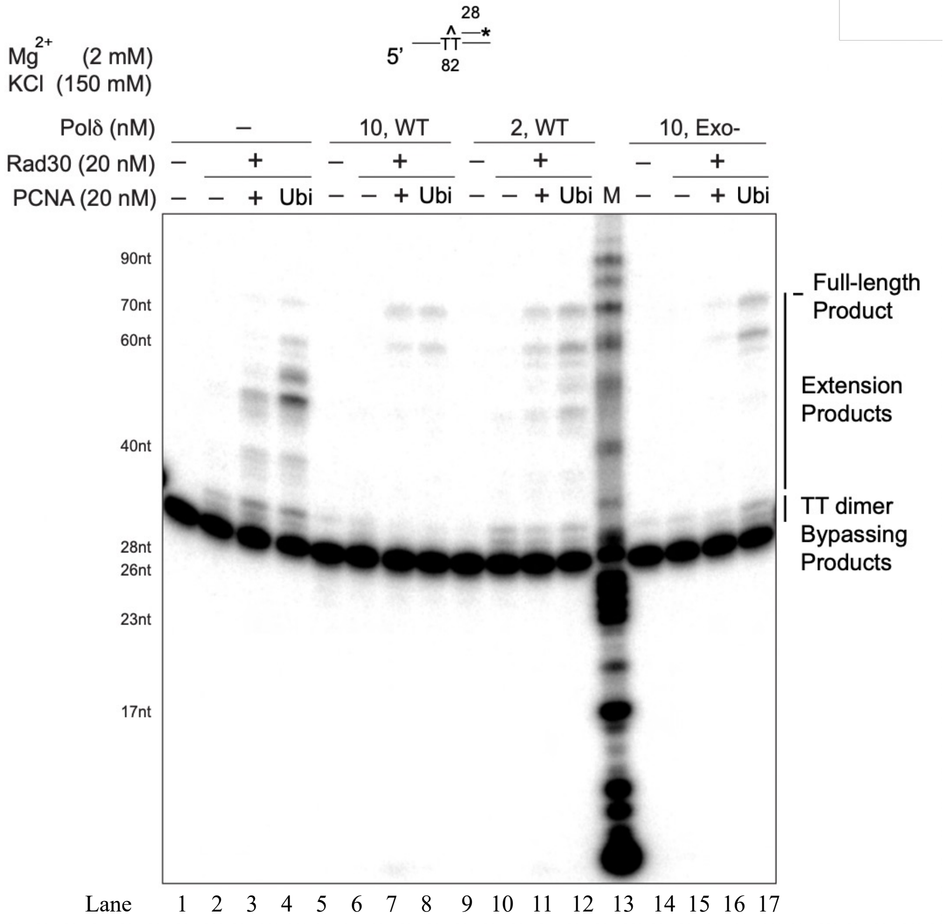

**Fig. S11. Pol δ physically blocks Rad30 accessing to the T-T dimer**. The stimulation of Rad30 lesion bypass by PCNA and ubiquitinylated-PCNA was tested under conditions of 2 mM $Mg^{2+}$ and 150 mM KCl in the absence or presence of Pol δ (10 nM and 2 nM) or the Pol δ exonuclease-deficient mutant.

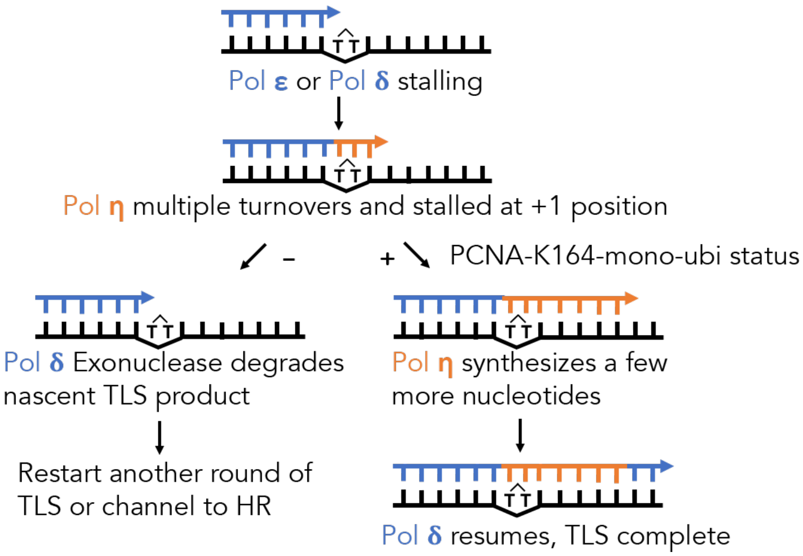

**Fig. S12. Model for the role of PCNA mono-ubiquitylation in TLS.** Without PCNA or Ub-PCNA, Rad30 could rarely finish TLS since it stalls at the first nucleotide downstream of a TT dimer. After PCNA is ubiquitylated by Rad18, TLS polymerases can proceed downstream of DNA lesion at further distance, escaping from the inhibition by Pol δ exonuclease. HR, homologous recombination.

**Table S1. List of yeast strains used in this study.**

| **Table S1. List of yeast strains used in this study** | | |
|---|---|---|
| Strain name | Genotype | Source |
| YB163 Mata | *MAT***a**-inc *ura3-52 his3*-Δ*200 ade2-101 lys2-801 trp1*-Δ*1 gal3– trp1::[his3*-Δ*3′::HOcs, his3*-Δ*5′]* | Fasullo M (*53*) |
| CY4865 | YB163 *pol3-5DV* | This study |
| CY4874 | YB163 *rad30Δ::Hyg* | This study |
| CY4907 | YB163 *pol3-5DV rad30Δ::Hyg* | This study |
| CY4970 | YB163 *rev3Δ::Kan* | This study |
| CY4905 | YB163 *pol3-5DV rev3Δ::Kan* | This study |
| CY4968 | YB163 *rev1Δ*::TRP1 | This study |
| CY5159 | *YB163 pol3-5DV rev1Δ::TRP1* | This study |
| CY5278 | *YB163 rev1Δ::TRP1 rev3Δ::Kan rad30Δ::Hyg* | This study |
| CY5281 | *YB163 pol3-5DV rad30Δ::Hyg rev3Δ::Kan rev1Δ:: TRP1* | This study |
| YKJM1 | *MAT***a** *ura3-52, leu2Δ1, trp1Δ63, his3Δ200, lys2ΔBgl, hom3-10, ade2Δ1, ade8, YEL069::URA3* | Myung KJ (*52*) |
| CY2657 | *YKJM1 rad14Δ::His3 clone1* | This study |
| CY3339 | YKJM1 *rad14Δ::His3 rev1Δ::Hyg* | This study |
| CY2678 | YKJM1 *rad14Δ::His3 rev3Δ::Kan* | This study |
| CY3658 | YKJM1 *rad14Δ::His3 rad30Δ::Nat* | This study |
| CY3514 | YKJM1 *rad14Δ::His3 rad30Δ::Nat rev1Δ::Hyg rev3Δ::Kan* (*tlsΔ*) | This study |
| CY4310 | YKJM1 *rad14Δ::His3 pol3-5DV* | This study |
| CY4481 | YKJM1 *rad14Δ::His3 pol3-5DV rev1Δ::Hyg* | This study |
| CY4573 | YKJM1 *rad14Δ::His3 pol3-5DV rev3Δ::Kan* | This study |
| CY4479 | YKJM1 *rad14Δ::His3 pol3-5DV rad30Δ::Nat* | This study |
| CY4444 | YKJM1 *rad14Δ::His3 rad30Δ::Nat rev1Δ::Hyg rev3Δ::Kan pol3-5DV* | This study |
| CY2743 | *YKJM1 rad14Δ::His3 rad51Δ::Kan* | This study |
| CY5375 | *YKJM1 rad14Δ::His3 rad51Δ::Kan pol3-5DV* | This study |
| CY4045 | *YKJM1 pol3-01* | This study |
| CY2630 | *YKJM1 siz1Δ::Kan* | This study |
| CY2554 | *YKJM1 ubc13Δ::Kan* | This study |
| CY4048 | *YKJM1 pol3-01 ubc13Δ::Kan* | This study |
| CY4073 | *YKJM1 ubc13Δ::Kan siz1Δ::NAT* | This study |
| CY4075 | *YKJM1 ubc13Δ::Kan siz1Δ::NAT pol3-01* | This study |
| CY2637 | *YKJM1 siz1Δ::Kan rad18Δ::His3* | This study |
| CY4080 | *YKJM1 pol3-01 rad18Δ::His3* | This study |
| CY4101 | *YKJM1 pol3-01 siz1Δ::NAT rad18Δ::His3* | This study |
| CY2609 | *YKJM1 rad51Δ::Kan* | This study |

| | | |
|---|---|---|
| CY2654 | *YKJM1 ubc13Δ::Kan rad51Δ::His3* | This study |
| CY4103 | *YKJM1 rad51Δ::Hyg pol3-01* | This study |
| CY4137 | *YKJM1 ubc13Δ::Kan rad51Δ::Hyg pol3-01* | This study |
| CY6657 | *YKJM1 rev3Δ::Kan* | This study |
| CY6661 | *YKJM1 rad30Δ::Hyg* | This study |
| CY5438 | *YKJM1 pol3-5DV* | This study |
| CY6765 | *YKJM1 rev3Δ::Kan pol3-5DV* | This study |
| CY6768 | *YKJM1 rad30Δ::Hyg pol3-5DV* | This study |
| CY2632 | *YKJM1 rad18Δ::His3* | This study |
| CY6568 | *YKJM1 rad18Δ::His3 pol3-5DV* | This study |
| CY6846 | *YKJM1 rad18Δ::His3 pol3-5DV rev3Δ::Kan* | This study |
| CY6843 | *YKJM1 rad18Δ::His3 pol3-5DV rad30Δ::Hyg* | This study |
| CY3102 | *YKJM1 pcnaK164R::LEU2* | This study |
| CY4333 | *YKJM1 pcnaK164R::LEU2 pol3-5DV clone1* | This study |
| CY8027 | *YKJM1 2HA-Rad30::LEU2* | This study |
| CY8030 | *YKJM1 rad18Δ::HIS3 2HA-Rad30::LEU2* | This study |
| CY8052 | *YKJM1 rad18Δ::HIS3 pol3-5DV 2HA-Rad30::LEU2* | This study |
| CY8067 | *YKJM1 2HA-rad30FFAA ::LEU2* | This study |
| CY8069 | *YKJM1 2HA-rad30-D570A ::LEU2* | This study |
| BY4741 | *MAT***a** *his3Δ1 leu2Δ0 met15Δ0 ura3Δ0* | |
| CY5026 | *BY4741 pol3-5DV* | This study |
| CY5013 | *BY4741 rad51Δ:::Hyg* | This study |
| CY5011 | *BY4741 ubc13Δ::Nat* | This study |
| CY5457 | *BY4741 ubc13::NAT pol3-5DV* | This study |
| CY5472 | *BY4741 ubc13Δ::Nat rad51Δ::Hyg* | This study |
| CY5813 | *BY4741 ubc13Δ::NAT rad51ΔKan pol3-5DV* | This study |
| CY6563 | *BY4741 rad14Δ::Leu2* | This study |
| CY6706 | *BY4741 rad14Δ::Leu2 pol3-5DV* | This study |
| CY8037 | *BY4741 rad14Δ::Leu2 pol3-5DV rad18Δ::His3* | This study |
| TGI354 | *hoΔ hml::ADE1 MAT***a***-inc hmr::ADE1 ade1 leu2-3,112 lys5 trp1::hisG ura3-52 ade3::GAL::HO arg5,6::MAT***a** | Ira G (*54*) |
| CY6703 | *TGI354 pol3-5DV* | This study |
| CY6963 | *TGI354 rad51Δ::Kan* | This study |
| CY6965 | *TGI354 rad51Δ::Kan pol3-5DV* | This study |
| W303-1A/α | *MAT***a** */MAT**α** {leu2-3,112 trp1-**1** can1-100 ura3-**1** ade2-**1** his3-11,15} [phi+]* | Zhao XL (*55*) |
| CY7646 | *W303-1 MAT**α** pol3-5DV* | This study |
| CY7713 | *W303-1 MAT**α** rev1Δ::Hyg rev3Δ::Kan rad30Δ::His3* | This study |

| | | |
|---|---|---|
| CY7717 | *W303-1 MAT***α** *rev1Δ::Hyg rev3Δ::Kan rad30Δ::His3 pol3-5DV* | This study |
| CY7721 | *W303-1 MAT***α** *rad51Δ::Ura3* | This study |
| CY7726 | *W303-1 MAT***α** *rad51Δ::Ura3 pol3-5DV* | This study |
| CY7730 | *W303-1 MAT***α** *rad51Δ::Ura3 rad30Δ::His3* | This study |
| CY7733 | *W303-1 MAT***α** *rad51Δ::Ura3 rad30Δ::His3 pol3-5DV* | This study |
| CY7736 | *W303-1 MAT***α** *rad51Δ::Ura3 rev1Δ::Hyg* | This study |
| CY7740 | *W303-1 MAT***α** *rad51Δ::Ura3 rev1Δ::Hyg pol3-5DV* | This study |
| CY7743 | *W303-1 MAT***α** *rad51Δ::Ura3 rev3Δ::Kan* | This study |
| CY7747 | *W303-1 MAT***α** *rad51Δ::Ura3 rev3Δ::Kan pol3-5DV* | This study |
| CY7751 | *W303-1 MAT***α** *rad51Δ::Ura3 rev1Δ::Hyg rev3Δ::Kan rad30Δ::His3* | This study |
| CY7755 | *W303-1 MAT***α** *rad51Δ::Ura3 rev1Δ::Hyg rev3Δ::Kan rad30Δ::His3 pol3-5DV* | This study |
| CY7768 | *W303-1 MAT***α** *rad52Δ::NAT* | This study |
| CY7774 | *W303-1 MAT***α** *rad52Δ::NAT pol3-5DV* | This study |
| CY7780 | *W303-1 MAT***α** *rad52Δ::NAT rev3Δ::Kan* | This study |
| CY7803 | *W303-1 MAT***α** *rad52Δ::NAT rev3Δ::Kan pol3-5DV* | This study |
| CY7807 | *W303-1 MAT***α** *rad52Δ::NAT rev1Δ::Hyg* | This study |
| CY7812 | *W303-1 MAT***α** *rad52Δ::NAT rev1Δ::Hyg pol3-5DV* | This study |
| CY7817 | *W303-1 MAT***α** *rad52Δ::NAT rad30Δ::His3* | This study |
| CY7822 | *W303-1 MAT***α** *rad52Δ::NAT rad30Δ::His3 pol3-5DV* | This study |
| CY7827 | *W303-1 MAT***α** *rad52Δ::NAT rev1Δ::Hyg rev3Δ::Kan rad30Δ::His3* | This study |
| CY7829 | *W303-1 MAT***a** *rad52Δ::NAT rev1Δ::Hyg rev3Δ::Kan rad30Δ::His3* | This study |
| CY7831 | *W303-1MAT***α** *rad52Δ::NAT rev1Δ::Hyg rev3Δ::Kan rad30Δ::His3 pol3-5DV* | This study |
| CY7833 | *W303-1 MAT***a** *rad52Δ::NAT rev1Δ::Hyg rev3Δ::Kan rad30Δ::His3 pol3-5DV* | This study |
| CY4703 | *W303-1 MAT***α** *pol3-01* | This study |
| CY8007 | *W303-1 MAT***α** *rad51Δ::Ura3 pol3-01* | This study |
| CY8014 | *W303-1 MAT***a** *rad51Δ::Ura3 pol3-01 rev1Δ::Hyg rev3Δ::Kan rad30Δ::His3* | This study |
| CY8011 | *W303-1 MAT***α** *rad52Δ::Nat pol3-01* | This study |
| CY8015 | *W303-1 MAT***a** *rad52Δ::Nat pol3-01 rev1Δ::Hyg rev3Δ::Kan rad30Δ::His3* | This study |
| CY4307 | *W303-1 MAT***a** *pol3-5DV* | This study |
| CY4370 | *W303-1 MAT***a** *rad18Δ::Nat* | This study |
| CY4380 | *W303-1 MAT***a** *rev1Δ::Hyg rev3Δ::KanMx rad30Δ::His3 rad18Δ::Nat* | This study |
| CY4421 | *W303-1 MAT***a** *rev1Δ::Hyg rev3Δ::KanMx rad30Δ::His3 rad18Δ::Nat pol3-5DV* | This study |
| CY4455 | *W303-1 MAT***a** *ubc13Δ::Kan clone1* | This study |
| CY4457 | *W303-1 MAT***a** *pol3-5DV ubc13Δ::Kan* | This study |
| CY4459 | *W303-1 MAT***a** *mms2Δ::Kan clone1* | This study |
| CY4461 | *W303-1 MAT***a** *pol3-5DV mms2Δ::Kan* | This study |

| | | |
|---|---|---|
| CY4530 | *W303-1 MAT***a** *rad5Δ::Kan clone1* | This study |
| CY4532 | *W303-1 MAT***a** *pol3-5DV rad5Δ::Kan* | This study |
| CY4434 | *W303-1 MAT***a** *rev1Δ::Hyg rev3Δ::KanMx rad30Δ::His3 pol3-5DV* | This study |
| CY4435 | *W303-1 MAT***a** *rev1Δ::Hyg rev3Δ::KanMx rad30Δ::His3 ubc13Δ::Nat* | This study |
| CY4436 | *W303-1 MAT***a** *rev1Δ::Hyg rev3Δ::KanMx rad30Δ::His3 ubc13Δ::Nat pol3-5DV* | This study |
| CY5165 | *W303-1 MAT***a** *mms2Δ::Kan rad30Δ His3 rev3Δ::Trp1 rev1Δ::Hyg* | This study |
| CY5168 | *W303-1 MAT***a** *mms2Δ::Kan rad30Δ His3 rev3Δ::Trp1 rev1Δ::Hyg pol3-5DV* | This study |
| CY5171 | *W303-1 MAT***a** *rad5Δ::Kan rad30Δ His3 rev3Δ::Trp1 rev1Δ::Hyg* | This study |
| CY5173 | *W303-1 MAT***a** *rad5Δ::Kan rad30Δ His3 rev3Δ::Trp1 rev1Δ::Hyg pol3-5DV* | This study |
| CY5266 | *W303-1 MAT***a** *pol2-04* | This study |
| CY5268 | *W303-1 MAT***a** *pol2-04 rad5Δ:: Kan* | This study |
| CY5270 | *W303-1 MAT***a** *pol2-04 ubc13Δ::Kan* | This study |
| CY5675 | *W303-1 MAT***a** *pol2-04 rad51Δ::Kan* | This study |
| CY5677 | *W303-1 MAT***a** *pol2-04 rad52Δ::Kan* | This study |
| CY5719 | *W303-1 MAT***a** *ubc13Δ::Kan siz1Δ::NAT* | This study |
| CY5722 | *W303-1 MAT***a** *ubc13Δ::Kan siz1Δ::NAT pol3-5DV* | This study |
| CY5977 | *W303-1 MAT***α** *pcnaK164R::LEU2* | This study |
| CY5979 | *W303-1 MAT***α** *pcnaK164R::LEU2 pol3-5DV* | This study |
| CY6005 | *W303-1 MAT***α** *pcnaK164R::LEU2 rev1Δ::Hyg rev3Δ::Kan rad30Δ::His3* | This study |
| CY6009 | *W303-1 MAT***α** *pcnaK164R::LEU2 rev1Δ::Hyg rev3Δ::Kan rad30Δ::His3 pol3-5DV* | This study |
| CY7704 | *W303-1 MAT***α** *rad30Δ::His3* | This study |
| CY7701 | *W303-1 MAT***α** *rad30Δ::His3 pol3-5DV* | This study |
| CY6749 | *W303-1 MAT***α** *rad30-D570A (thus ubz-)* | This study |
| CY6907 | *W303-1 MAT***a** *rad30-D570A pol3-5DV* | This study |
| CY7676 | *W303-1 MAT***α** *rad30-FFAA (thus, pip-)* | This study |
| CY7979 | *W303-1 MAT***α** *rad30-FFAA pol3-5DV* | This study |
| CY7631 | *w303 diploid WT\|WT* | This study |
| CY7634 | *w303 diploid rad18Δ::NAT\|rad18Δ::NAT* | This study |
| CY7632 | *w303 diploid rad18Δ::NAT pol3-5DV\|rad18Δ::NAT POl3-WT* | This study |
| CY7642 | *w303 diploid rad18Δ::NAT pol3-5DV\|rad18Δ::NAT pol3-5DV* | This study |

**Table S2. Spontaneous Forward CAN1 Mutation rate**

| Table S2. Spontaneous Forward CAN1 Mutation rate * | | |
|---|---|---|
| Strains | Rate($10^{-7}$) | Relative rate |
| *rad14Δ* | 4.82 ± 0.38 | 1.00 |
| *rad14Δ rev1Δ* | 0.81 ± 0.09 | 0.17 |
| *rad14Δ rev3Δ* | 0.76 ± 0.09 | 0.16 |
| *rad14Δ rad30Δ* | 4.88 ± 0.72 | 1.01 |
| *rad14Δ tlsΔ* | 1.08 ± 0.07 | 0.22 |
| *rad14Δ pol3-5DV* | 22.35 ± 1.35 | 4.64 |
| *rad14Δ pol3-5DV rev1Δ* | 24.80 ± 2.83 | 5.14 |
| *rad14Δ pol3-5DV rev3Δ* | 23.20 ± 2.97 | 4.81 |
| *rad14Δ pol3-5DV rad30Δ* | 22.45 ± 2.47 | 4.66 |
| *rad14Δ pol3-5DV tlsΔ* | 31.75 ± 1.20 | 6.59 |

* Mutation rate is determined by flutuation test with 7 or 9 individual clones for each strain and the experiments were repeated three times to get the standard deviation

**Table S3. The number of different type mutations found in WGS with or without 3J/m$^2$ UV treatment**

Table S3. The number of different type mutations found in WGS with or without 3J/m$^2$ UV treatment [1]

| Type of mutation[2] | *rad14Δ* no-UV | | *rad14Δ pol3-5DV* no-UV | | *rad14Δ* UV | | *rad14Δ 5DV* UV | | *rad14Δ 5DV rad18Δ* UV | | r14 5DV/r14 0J (fold) | r14 5DV/r14 3J (fold) |
|---|---|---|---|---|---|---|---|---|---|---|---|---|
| | No. | % | No. | % | No. | % | No. | % | No. | % | | |
| CC: 3' C > A | 0 | 0.00% | 0 | 0.00% | 2 | 0.03% | 5 | 0.04% | 2 | 0.02% | N.A.[3] | 2.50 |
| CC: 3' C > G | 1 | 0.29% | 9 | 2.15% | 3 | 0.04% | 140 | 1.17% | 392 | 2.99% | 9.00 | 46.67 |
| CC: 3' C > T | 1 | 0.29% | 2 | 0.48% | 50 | 0.67% | 227 | 1.89% | 61 | 0.47% | 2.00 | 4.54 |
| CC: 5' C > A | 1 | 0.29% | 1 | 0.24% | 35 | 0.47% | 78 | 0.65% | 18 | 0.14% | 1.00 | 2.23 |
| CC: 5' C > G | 6 | 1.75% | 9 | 2.15% | 9 | 0.12% | 617 | 5.15% | 1993 | 15.19% | 1.50 | 68.56 |
| CC: 5' C > T | 16 | 4.68% | 19 | 4.55% | 96 | 1.30% | 246 | 2.05% | 120 | 0.91% | 1.19 | 2.56 |
| CT:C > A | 0 | 0.00% | 0 | 0.00% | 1 | 0.01% | 4 | 0.03% | 10 | 0.08% | N.A. | 4.00 |
| CT:C > G | 8 | 2.34% | 12 | 2.87% | 7 | 0.09% | 417 | 3.48% | 958 | 7.30% | 1.50 | 59.57 |
| CT:C > T | 5 | 1.46% | 8 | 1.91% | 16 | 0.22% | 69 | 0.58% | 38 | 0.29% | 1.60 | 4.31 |
| CT:T > A | 5 | 1.46% | 7 | 1.67% | 49 | 0.66% | 71 | 0.59% | 38 | 0.29% | 1.40 | 1.45 |
| CT:T > C | 1 | 0.29% | 2 | 0.48% | 18 | 0.24% | 158 | 1.32% | 14 | 0.11% | 2.00 | 8.78 |
| CT:T > G | 0 | 0.00% | 0 | 0.00% | 3 | 0.04% | 2 | 0.02% | 1 | 0.01% | N.A. | 0.67 |
| TC:C > A | 0 | 0.00% | 1 | 0.24% | 2 | 0.03% | 8 | 0.07% | 12 | 0.09% | N.A. | 4.00 |
| TC:C > G | 16 | 4.68% | 12 | 2.87% | 9 | 0.12% | 300 | 2.50% | 956 | 7.29% | 0.75 | 33.33 |
| TC:C > T | 3 | 0.88% | 8 | 1.91% | 265 | 3.58% | 426 | 3.56% | 127 | 0.97% | 2.67 | 1.61 |
| TC:T > A | 57 | 16.67% | 61 | 14.59% | 2031 | 27.41% | 1516 | 12.65% | 1452 | 11.07% | 1.07 | 0.75 |
| TC:T > C | 12 | 3.51% | 13 | 3.11% | 127 | 1.71% | 195 | 1.63% | 155 | 1.18% | 1.08 | 1.54 |
| TC:T > G | 1 | 0.29% | 2 | 0.48% | 53 | 0.72% | 56 | 0.47% | 51 | 0.39% | 2.00 | 1.06 |
| TT: 3' T > A | 2 | 0.58% | 2 | 0.48% | 105 | 1.42% | 201 | 1.68% | 63 | 0.48% | 1.00 | 1.91 |
| TT: 3' T > C | 0 | 0.00% | 5 | 1.20% | 65 | 0.88% | 457 | 3.81% | 57 | 0.43% | N.A. | 7.03 |
| TT: 3' T > G | 3 | 0.88% | 3 | 0.72% | 14 | 0.19% | 20 | 0.17% | 9 | 0.07% | 1.00 | 1.43 |
| TT: 5' T > A | 19 | 5.56% | 15 | 3.59% | 926 | 12.50% | 924 | 7.71% | 286 | 2.18% | 0.79 | 1.00 |
| TT: 5' T > C | 0 | 0.00% | 2 | 0.48% | 24 | 0.32% | 50 | 0.42% | 12 | 0.09% | N.A. | 2.08 |
| TT: 5' T > G | 4 | 1.17% | 7 | 1.67% | 139 | 1.88% | 166 | 1.39% | 28 | 0.21% | 1.75 | 1.19 |
| Total SPY | 161 | 47.08% | 200 | 47.85% | 4049 | 54.64% | 6353 | 53.02% | 6853 | 52.25% | 1.24 | 1.57 |
| UPY | 122 | 35.67% | 139 | 33.25% | 2791 | 37.67% | 4223 | 35.24% | 4116 | 31.38% | 1.14 | 1.51 |
| NPY | 59 | 17.25% | 79 | 18.90% | 570 | 7.69% | 1406 | 11.73% | 2148 | 16.38% | 1.34 | 2.47 |
| Grand Total | 342 | 100.00% | 418 | 100.00% | 7410 | 100.00% | 11982 | 100.00% | 13117 | 100.00% | 1.22 | 1.62 |

[1] Without Chr.XII and Mitochondrial; 2. E.g. CC:3'C>A is an abbreviation of 5'-CC-3' dimer with its 3'C mutated to A; [3] N.A. Not applicable

### Table S4. The sequence of oligonucleotide used for biochemical analysis.

Table S4. The sequence of oligonucleotide used for biochemical analysis.

| NAME | SEQUENCE |
|---|---|
| H1 | CCCCCTCGAGGTCGACGGTATCGATAAGCTTGATATCGAA<u>TT</u>CCTGCAGCCCGGGGGATCCACTAGTTCTAGAGCGGCCGCC |
| H2 | CCCCCTCGAGGTCGACGGTATCGATAAGCTTGATATCGAA(T^T_dimer)CCTGCAGCCCGGGGGATCCACTAGTTCTAGAGCGGCCGCC |
| H3 | AGAACTAGTGGATCCCCCGGGCTGCA |
| H4 | AGAACTAGTGGATCCCCCGGGCTGCAGG |
| H5 | AGAACTAGTGGATCCCCCGGGCTGCAGGAA |
| H6 | AGAACTAGTGGATCCCCCGGGCTGCAGGAAT |
| H7 | AGAACTAGTGGATCCCCCGGGCTGCAGGAATT |
| H8 | AGAACTAGTGGATCCCCCGGGCTGCAGGAATTC |
| H9 | AGAACTAGTGGATCCCCCGGGCTGCAGGAATTCG |
| H10 | AGAACTAGTGGATCCCCCGGGCTGCAGGAATTCGAT |
| H11 | GGCGGCCGCTCTAGAACTAGTGGATCCCCCGGGCTGCAGGAATTCGATATCAAGCTTATCGATACCGTCGACCTCGAGGGGG |

**Data S1. SPY & NPY(no TC T>A&TT 5'T>A ) flanking efficient early replication origins (separate file).**

**Data S2. NPY-dPY pairs (IMD within 100 bp) (separate file).**
